# Functional signature of conversion in Mild Cognitive Impairment patients

**DOI:** 10.1101/290783

**Authors:** Stefano Delli Pizzi, Miriam Punzi, Stefano L Sensi, for the Alzheimer’s Disease Neuroimaging Initiative

**Affiliations:** Department of Neuroscience, Imaging and Clinical Sciences, “G. d’Annunzio” University, Chieti, Italy; Center for excellence on Aging and Translational Medicine - Ce.S.I. - Me.T., “G. d’Annunzio” University, Chieti, Italy; Departments of Neurology and Pharmacology, Institute for Memory Impairments and Neurological Disorders, University of California-Irvine, Irvine, CA, USA.

**Keywords:** Alzheimer’s Disease, entorhinal cortex, functional connectivity, Hippocampus, Mild Cognitive Impairment

## Abstract

The entorhinal-hippocampal circuit is a strategic hub for memory but also the first site to be affected in the Alzheimer’s Disease (AD)-related pathology. We investigated MRI patterns of brain atrophy and functional connectivity in a study cohort obtained from the Alzheimer’s Disease Neuroimaging Initiative database including healthy control (HC), Mild Cognitive Impairment (MCI), and AD subjects. MCI individuals were clinically evaluated 24 months after the MRI scan, and the group further divided into a subset of subjects who either did (c-MCI) or did not (nc-MCI) convert to AD. Compared to HC subjects, AD patients exhibited a collapse of long-range connectivity from the hippocampus and entorhinal cortex, pronounced cortical/sub-cortical atrophy, and a dramatic decline in cognitive performances. c-MCI patients showed entorhinal and hippocampal hypo-connectivity, no signs of cortical thinning but evidence of right hippocampus atrophy. On the contrary, nc-MCI patients showed lack of brain atrophy, largely preserved cognitive functions, hippocampal and entorhinal hyper-connectivity with selected neocortical/sub-cortical regions mainly involved in memory processing and brain meta-stability. This hyper-connectivity can represent an early compensatory strategy to overcome the progression of cognitive impairment. This functional signature can also be employed for the diagnosis of c-MCI subjects.

## Introduction

Brain aging and aging-related neurodegenerative disorders are a significant health challenge for contemporary societies. Brain aging represents a favorable background for the onset and development of neurodegeneration and dementia. Alzheimer’s Disease (AD) is a condition associated with the development of irreversible cognitive and behavioral deficits and preceded by a prodromal stage known as Mild Cognitive Impairment (MCI). MCI patients do not fulfill the diagnostic criteria for dementia but show significant cognitive deficits that mostly occur in mnemonic domains (Petersen et al. 2010). The MCI stage progresses to AD in 60-65% of cases (Busse et al. 2006) with a conversion rate that reaches 8.1% per year (Mitchell and Shiri-Feshki 2009). Thus, the early identification of the brain changes associated with MCI is critical to catch the disease at its initial stage, unravel the pathogenic mechanisms involved in AD and help the design of more effective therapeutic interventions.

Neuroimaging approaches have been extensively employed to detect the initial changes associated with the early stages of AD (Frisoni and Jessen 2018). Resting-state functional Magnetic Resonance Imaging (rs-fMRI) is a non-invasive tool that allows the investigation of the operational changes and network reconfigurations that occur in several neurological/neurodegenerative conditions including AD. In MCI patients, this technique has been successfully employed in the quest to detect abnormalities in the brain functional connectivity that occur before the appearance of patent signs of structural damage (Badhwar et al. 2017, Drzezga et al. 2011, Canuet et al. 2015).

The entorhinal-hippocampal circuit is a strategic region for the control of cognitive processes and the first site to be affected by the AD-related pathology (Braak et al. 2013, Gomez-Isla et al. 1996). In the AD brain, the early signs of synaptic degradation occur within the perforant path, the neurodegeneration then spreads to the layers II-III of the entorhinal cortex and the hippocampal CA3/DG regions, eventually reaches the subicular areas, and ultimately affects the whole hippocampus (Yassa et al. 2010). The entorhinal-hippocampal complex plays a critical role in the processing of long-term memory (Preston and Eichenbaum 2013). The region sustains the network brain stability and promotes the adaptive neuroplasticity that copes with the underlying pathological stressors that are triggering the structural damage (van den Heuvel and Sporns 2011, Hillary and Grafman 2017).

In this study, we investigated, in a cohort of one hundred thirty-five individuals, differences in structural MRI (sMRI) and rs-fMRI features that occurred within the cortico-hippocampal and cortico-entorhinal circuits. The study group included Healthy Control (HC) (n=40), MCI (n=67), and AD (n=28) subjects. The dataset also provided information on the demographic, neuropsychological/clinical, and APOE status as well as the CSF levels of AD-related pathogenic proteins like the amyloid β_1–42_ peptide (Aβ_1–42_), tau phosphorylated at threonine 181 (p-tau_181_), and the ratio of p-tau_181/_Aβ_1–42_.

sMRI data were employed to investigate differences in brain volume and cortical thickness among study participants. Rs-fMRI data were used to evaluate differences in the functional connectivity (FC) occurring in the circuits linking the hippocampus or the entorhinal cortex to the cortex. The progression or clinical stability of HC or MCI subjects was assessed by using clinical follow-up data obtained 24 months after the initial MRI session. With the help of these longitudinal data, the MCI group was therefore divided into two subsets: patients who converted (c-MCI) or did not convert (nc-MCI) to AD. Finally, the CSF data were plotted against the rs-fMRI results to explore correlations between the FC strength and levels of Aβ_1–42_ and p-tau_181_ as well as the p-tau_181_/ Aβ_1–42_ ratio.

The overall aim of the study was to disclose the contributions of the hippocampus and entorhinal cortex in ongoing neurodegenerative processes and the transition from different steps of the AD-related spectrum.

## Materials and methods

### Experimental design

Data employed for this article were obtained from the ADNI-GO/2 database. ADNI was launched in 2003 as a public-private partnership led by Michael W. Weiner. The primary goal of ADNI is to employ serial MRI, PET, biological markers, and clinical and neuropsychological data to investigate the features of patients affected by the AD spectrum. For up-to-date information on the initiative, see www.adni-info.org.

Experiments fulfilled the ethical standards and the Declaration of Helsinki (1997) and subsequent revisions. Informed consent was obtained from study participants or authorized representatives. Study participants had a good general health status and no diseases that are expected to interfere with the study. Overall, the ADNI-GO/2 database included one hundred seventy participants who have completed the 3T-sMRI and 3T-rs-fMRI and with an age range between 57 and 88 years old.

Participants who did not complete a clinical follow-up performed 24 months after the first MRI session or those who showed technical issues related to their MRI raw-data (i.e., artifacts, dishomogeneity in acquisition parameters, images deformed for missing information raw-file) were excluded from the study sample (**Supplementary Fig. 1**). Our final sample included one hundred thirty-five participants divided into forty HC subjects, sixty-seven MCI patients, and twenty-eight AD patients. Based on the clinical follow-up, the MCI group was further subdivided into a group of fifty-four nc-MCI patients and thirteen c-MCI patients.

### Neuropsychological assessment

All subjects underwent clinical and cognitive evaluations at the time of the MRI scan. The ADNI neuropsychological dataset includes the Mini-Mental State Examination (MMSE) (Folstein et al. 1975) and the Montreal Cognitive Assessment (MoCA) (Nasreddine et al. 2005) to investigate global cognition; the Functional Activities Questionnaire (FAQ) for the assessment of daily living activities (Pfeffer et al. 1982); the Alzheimer’s Disease Assessment Scale-Cognitive subscales (ADAS - 11 items scores; ADAS - 13 items scores) to evaluate the severity of impairments of memory, learning, language (production and comprehension), praxis, and orientation (Mohs and Cohen 1988; Mohs et al. 1997); the Animal Fluency (Morris et al. 1989) and the 30-item Boston Naming Test (BNT) (Kaplan, et al. 1983) to investigate semantic memory and language abilities; the Trail Making Test (TMT), part A and B (time to completion) to assess attention/executive functions (Spreen 1998); the Rey Auditory Verbal Learning Test (RAVLT) to investigate recall and recognition (Rey, 1964).

HC subjects were free of memory complaints and without significant impairment as far as general cognitive functions or daily living activities. The inclusion criteria for HC subjects were: MMSE scores between 24 and 30, a global score of 0 on the Clinical Dementia Rating Scale (CDR-RS; Morris JC 1993), and a score above the cutoff level on the Logical Memory II, subscale of the Wechsler Memory Scale-Revised (WMS-R; Wechsler, 1987) (≥ for 0-7 years of education, ≥ for 8-15 years, and ≥ for 16 or more years).

The inclusion criteria for MCI patients were: MMSE scores between 24 and 30, memory impairments identified by the partner with or without complaints by the participant, a CDR score of 0.5, and memory deficits as indicated by scores below the cutoff level on the WMS-R Logical Memory II (0-7 for years of education 3-6, 5-9 for 8-15 years, and for >15 years ≤ Their general cognition status and functional performances were sufficiently preserved to exclude a diagnosis of AD.

AD patients fulfilled the criteria of probable AD, set by the National Institute of Neurologic and Communicative Disorders and Stroke (NINCDS) as well as the Alzheimer’s Disease and Related Disorders Association (ADRDA). AD patients had MMSE scores between 20 and 26 and CDR-RS scores between 0.5 and 1.0.

### CSF and APOE genotyping

CSF data were available for 87.4% of the total sample. The set included information on levels of the amyloid β_1–42_ peptide (Aβ_1–42_), total tau (t-tau), and tau phosphorylated at threonine 181 (p-tau_181_). Highly standardized Aβ_1–42_, t-tau and p-tau_181_ levels were measured using the Roche automated immunoassay platform (Cobas e601) and immunoassay reagents. Details on the methods for the acquisition and measurement of CSF are reported at the ADNI website (http://www.adni-info.org). The apolipoprotein E (APOE) ε allele frequency was also investigated at the screening stage.

### MRI acquisition protocol

MR data were acquired with a Philips 3T scanner (see details at http://adni.loni.usc.edu/wp-content/uploads/2010/05/ADNI2_MRI_Training_Manual_FINAL.pdf). T_1_-weighted images were obtained using 3D Turbo Field-Echo sequences (TFE, Slice Thickness=1.2 mm; TR/TE=6.8/3.1 ms). One run of Resting-state Blood Oxygen Level Dependent (BOLD) fMRI data was acquired using gradient-echo T_2_*-weighted echo-planar (EPI) sequence (in-plane voxel size=3.3125 mm x 3.3125 mm, slice thickness 3.3125 mm, and TR/TE=3000/30 ms. Subjects were instructed to lay still and keep their eyes open during acquisition.

### MRI data analysis

FreeSurfer (version 6.0) was employed to perform sMRI and rs-fMRI data analysis. For each study participants, T_1_-weighted images were analyzed using the “recon-all -all” command line to obtain automated reconstruction and labeling of cortical and subcortical regions (Fischl et al. 2004). The pre-processing steps encompassed magnetic field inhomogeneity correction, affine-registration to Talairach Atlas, intensity normalization and skull-strip. The processing steps involved segmentation of the subcortical white matter (WM) and deep grey matter (GM) volumetric regions, tessellation of the GM and WM matter boundary, automated topology correction, surface deformation following intensity gradients to optimally position the GM and WM and GM/cerebrospinal fluid borders at the location where the greatest shift in intensity delineates the transition to the other tissue class. The total volume of the left and right hippocampi and the estimated total intracranial volume (eTIV) were calculated using the “asegstats2table”. The hippocampal volumes were normalized by eTIV. The left and right masks of the hippocampi and entorhinal cortices were obtained by the “recon-all -all” command lines and used as “seed regions” for FC analysis using FreeSurfer - Functional Analysis Stream (http://surfer.nmr.mgh.harvard.edu/fswiki/FsFastFunctionalConnectivityWalkthrough). The ‘‘preprocsess’’ command line was employed to perform motion and slice timing corrections, masking, registration to the structural image, sampling to the surface, and surface smoothing by 5 mm as well as sampling to the MNI305 with volume smoothing. Surface sampling of time-series data was carried out onto the surface of the left and right hemispheres of the “fsaverage” template of FreeSurfer. Nuisance regressors were obtained for each study participants by extracting the EPI average time courses within the ventricle mask and the white matter mask (taking into consideration the top 5 principal components). These regressors, the motion correction parameters, and a fifth order polynomial were eliminated from the EPI time series. Temporal band-pass filtering (0.01<Hz<0.1) was applied to analyze only rs-fMRI data within this frequency range. The first four rs-fMRI time points were discarded to allow T_1_-weighted equilibration of the MRI signal. The mean signal time course within each seed region was employed as “regressor” to assess FC. With the “selxavg3-sess” command line, we performed the first level analysis (single subject analysis) including the computation of the Pearson correlation coefficient (r-value) between the time series within the seed and the time series at each voxel. The obtained correlation maps were then converted to Z-score maps before entering the second level analysis (group analysis). The “isxconcat-sess” command line was employed to create a “stack” of maps from each subject.

The Desikan-Killiany’s Atlas (Desikan et al. 2006) was employed to identify the location of clusters displaying structural MRI differences. In addition, two functional atlases, focused on cortical (Yeo et al. 2011) and cerebellar (Buckner et al. 2011) networks, were used to integrate the information provided by the Desikan-Killiany’s Atlas and define the positioning of clusters showing between-group differences or within-group correlations.

### Statistical analysis

One-way analysis of variance (ANOVA) and Bonferroni post-hoc test were employed to evaluate the group differences regarding demographic/clinical data as well as the hippocampal and entorhinal morphometry. Chi-squared test was used to investigate the group differences on gender and the APOE ε4 carrier status. For analyses related to FC and cortical thickness, general linear models (https://surfer.nmr.mgh.harvard.edu/fswiki/FsgdFormat) were used. The analysis investigated the differences between groups (I comparison: HC subjects, MCI patients, AD patients; II comparison: HC subjects, nc-MCI patients, c-MCI patients, AD patients). Moreover, further general linear models were used, in the MCI patients, to assess, relationships between FC strength and other variables of interest like the subject age, the seed morphometric measures, the cortical thickness in each vertex, and the CSF biomarkers. The correlation analyses between the variables of interest and the FC of a given seed region FC were performed in a vertex-by-vertex computation by using the “pvr” option in “mri_glmfit”. Using the “mri_concat” command line, conjunction maps were created to highlight the sites of overlaps occurring between clusters expressing significant group difference (HC vs. MCI) and clusters that indicate significant correlations between FC strength and variables of interest. All the results are shown on statistical maps and adjusted by applying cluster-wise correction for multiple comparisons (Hagler et al. 2006).

## Results

### Demographic and clinical features of the study groups

Global cognition and episodic memory (recall and recognition) were affected in MCI and AD patients. When compared to MCI and HC subjects, AD patients showed significant impairment of semantic memory, verbal fluency, language ability, and executive functions. AD patients also showed significantly higher frequency of APOE ε4 when compared to MCI or HC subjects. Within the MCI subsets, the global cognition and episodic memory (recall) were found to be more compromised in the c-MCI group. Between the two MCI subsets, no differences were found when considering other neuropsychological and clinical features or the APOE ε4 frequency. When compared to HC, levels of Aβ_1–42_, t-tau and p-tau_181_ were found to be altered in AD patients. No statistically significant differences were found when comparing levels of CSF biomarkers in HC vs. nc-MCI or c-MCI vs AD. Higher levels of t-tau, p-tau_181_, Aβ_1–42_/t-tau, and p-tau_181_/ Aβ_1–42_ as well as lower concentrations of Aβ_1–42_ were found in c-MCI patients when compared to HC subjects, and in AD patients when compared to both the nc-MCI and HC subjects. No differences regarding age and educational levels were observed among the study groups (HC, MCI, and AD) or the MCI subsets (nc-MCI and c-MCI). Statistics on demographics and clinical features of the study groups are shown in **Tables 1** and **2**.

**Table 1.** Demographic, neuropsychological and clinical features - I model

|  | HC | MCI | AD | ANOVA<sup>d</sup> |  | Post-hoc |  |  |
| --- | --- | --- | --- | --- | --- | --- | --- | --- |
| | | | | F or $\mu_2$ | p-value | HC vs. MCI | MCI vs. AD | HC vs. AD |
| N (% Male) | 40 (28%) | 67 (57%) | 28 (43%) | 8.709 | **0.013** | **0.003** | 0.217 | 0.188 |
| Age (y) | 74.1±5.7 | 71.6±6.9 | 72.3±6.8 | 0.709 | 0.494 | NA | NA | NA |
| Education (y) | 16.6±2.6 | 15.9±2.6 | 15.4±2.6 | 2.268 | 0.108 | NA | NA | NA |
| CDR-RS | 0.0±0.0 | 1.54±0.99 | 4.52±1.36 | 162.247 | **<0.001** | **<0.001** | **<0.001** | **<0.001** |
| MMSE | 29.0±1.4 | 27.7±1.7 | 22.6±2.6 | 80.871 | **<0.001** | **0.012** | **<0.001** | **<0.001** |
| MOCA | 25.6±2.2 | 23.0±3.0 | 15.6±4.9 | 62.383 | **<0.001** | **0.006** | **<0.001** | **<0.001** |
| LM | 13.6±3.1 | 7.1±3.0 | 1.3±2.0 | 131.622 | **<0.001** | **<0.001** | **<0.001** | **<0.001** |
| ADAS11 | 6.1±2.7 | 9.4±3.9 | 23.5±8.2 | 101.833 | **<0.001** | **0.004** | **<0.001** | **<0.001** |
| ADAS13 | 9.5±4.1 | 15.0±5.8 | 34.6±9.6 | 118.671 | **<0.001** | **<0.001** | **<0.001** | **<0.001** |
| FAQ | 0.1±0.5 | 3.2±4.1 | 16.1±7.2 | 98.961 | **<0.001** | **0.002** | **<0.001** | **<0.001** |
| AF | 20.7±4.6 | 18.6±4.8 | 12.8±5.1 | 26.437 | **<0.001** | 0.065 | **<0.001** | **<0.001** |
| BNT | 28.5±1.4 | 26.8±3.5 | 22.7±6.1 | 19.625 | **<0.001** | 0.154 | **<0.001** | **<0.001** |
| TMT-A<sup>#</sup> | 35.7±13.4 | 38.9±15.6 | 70.1±30.5 | 36.494 | **<0.001** | 1.000 | **<0.001** | **<0.001** |
| TMT-B | 87.7±55.6 | 113.1±67.1 | 235.2±81.5 | 47.062 | **<0.001** | 0.260 | **<0.001** | **<0.001** |
| RAVLT-IR<sup>##</sup> | 7.1±3.5 | 3.9±3.6 | 0.3±1.1 | 29.479 | **<0.001** | **<0.001** | **<0.001** | **<0.001** |
| RAVLT-RN<sup>#</sup> | 13.1±2.0 | 10.8±2.9 | 5.8±4.0 | 37.479 | **<0.001** | **0.001** | **<0.001** | **<0.001** |
| APOE ε4 | 33% | 42% | 75% | 12.895 | **0.002** | 0.289 | **0.010** | **0.001** |
| Aβ (pg/ml)<sup>###</sup> | 1375±651 | 1132±594 | 740±392 | 8.543 | **<0.001** | 0.159 | **0.018** | **<0.001** |
| tau (pg/ml)<sup>###</sup> | 246±68 | 298±137 | 368±152 | 6.525 | **0.002** | 0.170 | 0.076 | **0.001** |
| p-tau (pg/ml)<sup>###</sup> | 23±7 | 28±15 | 35±15 | 5.795 | **0.004** | 0.187 | 0.122 | **0.003** |
| t-tau/Aβ<sup>###</sup> | 0.23±0.17 | 0.37±0.34 | 0.58±0.31 | 10.039 | **<0.001** | 0.096 | **0.010** | **<0.001** |
| p-tau/Aβ<sup>###</sup> | 0.03±0.03 | 0.04±0.04 | 0.056±0.030 | 5.849 | **0.004** | 0.582 | **0.035** | **0.003** |
| L-HP/eTIV<sup>a,b</sup> | 2.58±0.30 | 2.44±0.44 | 2.01±0.40 | 22.370 | **<0.001** | 0.064 | **<0.001** | **<0.001** |
| R-HP/eTIV<sup>a,b</sup> | 2.64±0.34 | 2.49±0.49 | 2.11±0.43 | 18.301 | **<0.001** | 0.085 | **<0.001** | **<0.001** |
| eTIV<sup>c</sup> | 1.49±0.21 | 1.52±0.21 | 1.52±0.24 | 0.467 | 0.628 | NA | NA | NA |

Values are expressed as the mean ± standard deviation (SD). <sup>#</sup> Data on TMA-A were available for 134/135
participants (data was missing for one AD patient). <sup>##</sup> Data on RAVLT-DR and RAVLT-RE were available for
134/135 participants (RAVLT scores were missing for one MCI patient). <sup>###</sup> Data on CSF biomarkers were available
for 118/135 participants (34 HC subjects, 59 MCI patients, 25 AD patients). <sup>a</sup> Right (R) and left (L) hippocampus
(HP) volume is normalized for estimated total intracranial volume (eTIV); <sup>b</sup> values $\times 10^{-3}$ ; <sup>c</sup> values $\times 10^6$ ; <sup>d</sup> I model:
comparison between HC, patients with MCI and patients with AD.
*Abbreviations:* ADAS=Alzheimer's Disease Assessment Scale; AVL=Rey Auditory Verbal Learning Test;
BNT=Boston Naming Test; CDR-RS=Clinical Dementia Rating Scale; MMSE=Mini-Mental State Examination;
MoCA=Montreal Cognitive Assessment; TMT=Trail Making Test; $A\beta_{1-42}$ =amyloid $\beta_{1-42}$ ; p-tau<sub>181</sub>=tau phosphorylated at threonine 181; RAVLT-DR=Rey's Auditory Verbal Learning Test, 30 minute Delayed Recall; RAVLT-RN=Rey's Auditory Verbal Learning Test Delayed Recognition; eTIV=estimated total intracranial volume; EC=entorhinal cortex; HP=hippocampus;

**Table 2.** Demographic, neuropsychological and clinical features - II model

|  | nc-MCI | c-MCI | ANOVA-II model<sup>d</sup> |  | Post-hoc |  |  |  |  |  |
| --- | --- | --- | --- | --- | --- | --- | --- | --- | --- | --- |
| | | | F or $\mu_2$ | p-value | HC vs.<br>nc-MCI | HC vs.<br>c-MCI | nc-MCI<br>vs. c-MCI | nc-MCI<br>vs. AD | c-MCI<br>vs. AD | HC vs.<br>AD |
| N (% Male) | 54 (54%) | 13 (69%) | 8.779 | **0.032** | **0.011** | **0.007** | 0.310 | 0.352 | 0.116 | 0.188 |
| Age (y) | 71.6±7.1 | 71.7±6.1 | 0.470 | 0.703 | NA | NA | NA | NA | NA | NA |
| Education (y) | 15.9±2.7 | 16.2±2.5 | 1.558 | 0.203 | NA | NA | NA | NA | NA | NA |
| CDR-RS | 1.2±0.7 | 2.8±1.0 | 150.878 | **<0.001** | **<0.001** | **<0.001** | **<0.001** | **<0.001** | **0.012** | **<0.001** |
| MMSE | 27.9±1.7 | 27.0±1.6 | 55.520 | **<0.001** | **0.129** | **0.012** | 0.667 | **<0.001** | **<0.001** | **<0.001** |
| MOCA | 23.1±3.2 | 22.6±2.5 | 41.567 | **<0.001** | **0.038** | 0.062 | 1.000 | **<0.001** | **<0.001** | **<0.001** |
| LM | 7.2±3.0 | 6.5±3.0 | 87.650 | **<0.001** | **<0.001** | **<0.001** | 1.000 | **<0.001** | **<0.001** | **<0.001** |
| ADAS11 | 8.7±3.5 | 12.2±4.3 | 72.559 | **<0.001** | 0.084 | **0.001** | 0.114 | **<0.001** | **<0.001** | **<0.001** |
| ADAS13 | 14.0±5.2 | 19.1±6.2 | 86.011 | **<0.001** | **0.013** | **<0.001** | **0.046** | **<0.001** | **<0.001** | **<0.001** |
| FAQ | 2.1±3.4 | 8.0±3.1 | 84.147 | **<0.001** | 0.157 | **<0.001** | **<0.001** | **<0.001** | **<0.001** | **<0.001** |
| AF | 18.7±4.8 | 17.8±4.5 | 17.733 | **<0.001** | 0.295 | 0.235 | 1.000 | **<0.001** | **0.004** | **<0.001** |
| BNT | 27.1±3.1 | 25.8±4.9 | 13.471 | **<0.001** | 0.740 | 0.229 | 1.000 | **<0.001** | **0.041** | **<0.001** |
| TMT-A<sup>#</sup> | 37.7±14.9 | 44.0±18.0 | 25.197 | **<0.001** | 1.000 | 0.587 | 0.975 | **<0.001** | **<0.001** | **<0.001** |
| TMT-B | 105.5±60.4 | 144.4±85.5 | 34.944 | **<0.001** | 1.000 | **0.017** | 0.080 | **<0.001** | **<0.001** | **<0.001** |
| RAVLT-IR<sup>##</sup> | 4.5±3.7 | 1.5±1.9 | 23.883 | **<0.001** | **0.002** | **<0.001** | **0.023** | **<0.001** | 1.000 | **<0.001** |
| RAVLT-RN<sup>#</sup> | 11.0±3.0 | 9.7±2.3 | 25.805 | **<0.001** | **0.018** | **0.005** | 1.000 | **<0.001** | **0.005** | **<0.001** |
| APOE ε4 | 37% | 62% | 15.428 | **0.001** | 0.415 | 0.082 | 0.274 | **0.004** | 0.622 | **0.001** |
| Aβ (pg/ml)<sup>###</sup> | 1246±601 | 727±357 | 8.976 | **<0.001** | 1.000 | **0.003** | **0.023** | **0.003** | 1.000 | **0.002** |
| tau (pg/ml)<sup>###</sup> | 273±119 | 389±160 | 7.757 | **<0.001** | 1.000 | **0.003** | **0.017** | **0.015** | 1.000 | **0.002** |
| p-tau (pg/ml)<sup>###</sup> | 25.4±12.5 | 39±19 | 8.294 | **<0.001** | 1.000 | **0.001** | **0.004** | **0.017** | 1.000 | **0.003** |
| t-tau/Aβ<sup>###</sup> | 0.29±0.24 | 0.65±0.47 | 13.494 | **<0.001** | 1.000 | **<0.001** | **0.001** | **0.001** | 1.000 | **<0.001** |
| p-tau/Aβ<sup>###</sup> | 0.03±0.02 | 0.07±0.05 | 9.843 | **<0.001** | 1.000 | **0.001** | **0.003** | **0.002** | 1.000 | **0.004** |
| L-HP/eTIV<sup>a,b</sup> | 2.47±0.44 | 2.28 ±0.42 | 15.710 | **<0.001** | 0.449 | 0.062 | 0.961 | **<0.001** | 0.056 | **<0.001** |
| R-HP/eTIV<sup>a,b</sup> | 2.56±0.46 | 2.23±0.53 | 14.620 | **<0.001** | 0.940 | **0.008** | 0.110 | **<0.001** | 0.679 | **<0.001** |
| eTIV<sup>c</sup> | 1.51±0.21 | 1.56±0.23 | 0.442 | 0.723 | NA | NA | NA | NA | NA | NA |

Values are expressed as the mean ± standard deviation (SD). <sup>#</sup>TMA-A score was missing for one Alzheimer's Disease (AD) patient. <sup>##</sup>RAVLT scores were missing for one nc-MCI patient. <sup>###</sup>Data on CSF biomarkers were available for 51 nc-MCI patients and seven c-MCI patients. <sup>a</sup> right (R) and left (L) hippocampus (HP) volume is normalized for estimated total intracranial volume (eTIV); <sup>b</sup> values $\times 10^{-3}$ ; <sup>c</sup> values $\times 10^6$ ; <sup>d</sup> II model: comparison between HC, nc-MCI, c-MCI, AD.
*Abbreviations:* ADAS=Alzheimer's Disease Assessment Scale; AVLT=Rey Auditory Verbal Learning Test;
BNT=Boston Naming Test; CDR-RS=Clinical Dementia Rating Scale; MMSE=Mini-Mental State Examination;
MoCA=Montreal Cognitive Assessment; TMT=Trail Making Test; $A\beta_{1-42}$ =amyloid $\beta_{1-42}$ ; p-tau<sub>181</sub>=tau phosphorylated at threonine 181; RAVLT-DR=Rey's Auditory Verbal Learning Test, 30 minute Delayed Recall; RAVLT-RN=Rey's Auditory Verbal Learning Test Delayed Recognition; eTIV=estimated total intracranial volume; EC=entorhinal cortex; HP=hippocampus;

### Morphometric variations in the study groups

AD patients showed hippocampal atrophy. No statistically significant differences in hippocampal volumes were found between the two MCI subgroups. However, when compared to HC subjects, c-MCI exhibited signs of atrophy in the right hippocampus. AD patients showed bilateral hippocampal atrophy when compared to HC or nc-MCI patients whereas no differences in hippocampal volumes were found when compared to c-MCI patients. No differences in the estimated total intracranial volume (eTIV) were seen in the study groups or the MCI subgroups. The statistical analysis of the structural data is shown in **Tables 1** and **2**.

Cortical thinning was found in AD patients when their cortical thickness was compared to HC (**Supplementary Fig. 2A; Supplementary Fig. 3A**) or MCI subjects (**Supplementary Fig. 2B**). The cortical atrophy, found in AD patients, was more prominent in the insula, the temporo-occipital areas, the supramarginal and angular, the temporo-parietal junction, the posterior cingulate cortex/precuneus, and the parahippocampal and entorhinal cortices. These regions actively participate in the connectivity patterns of the Default-Mode Network (DMN) and the Posterior Medial Network (PMN), thereby indicating the presence of disease-driven differences of critical functional value. No significant differences in cortical thinning were instead observed when comparing MCI vs. HC subjects.

In AD patients, the comparison with nc-MCI (**Supplementary Fig. 3A**) or HC (**Supplementary Fig. 3C**) subjects indicated greater thinning of brain regions that belong to the DMN and PMN. Significant thinning of the mesial temporal regions was instead found in AD patients when compared to c-MCI patients (**Supplementary Fig. 3B**). Finally, no statistically significant differences in cortical thickness were instead found when comparing the c-MCI with nc-MCI patients or HC subjects.

### The entorhinal and hippocampal FC shows a distinct pattern in MCI subjects compared to AD patients

The analysis of rs-fMRI data revealed the presence of increased hippocampal FC in the MCI patients while the FC was decreased in AD patients. The investigation of the regional distribution of FC differences indicated that, compared to HC subjects (**Supplementary Fig. 4A**), MCI individuals exhibited enhanced connectivity of the hippocampus with the right medial prefrontal cortex (mPFC), cerebellar regions that are part of the DMN, right hypothalamus and left caudate, and hypo-connectivity between the hippocampus and of cerebellar regions that are part of the SN/CON. On the other hand, compared to MCI (**Supplementary Fig. 4B; Fig. 3B**) or HC (**Supplementary Fig. 4C**) individuals, AD subjects showed a decreased FC that took place in the DMN/PMN, striatum, and brainstem, and cerebellum regions that are part of the sensorimotor network. The entorhinal FC was found to be unaffected in MCI subjects and reduced in AD patients. Compared to HC subjects, AD patients displayed hypo-connectivity with DMN-related regions as well as with brainstem, striatum and cerebellar areas that are functionally linked to attentional and sensorimotor networks (**Supplementary Fig. 5B**, for the I model; **Fig. 4C, Supplementary Table 1**, for the II model). In addition, compared to MCI subjects, AD patients showed hypo-connectivity with cerebellar areas as well as with brainstem and thalamus (**Supplementary Fig. 5A**).

### Divergent patterns of functional connectivity within the MCI subgroups

The analysis of the hippocampal FC of the two MCI groups identified significant differences. Nc-MCI patients were characterized by enhanced FC of the hippocampus with the mPFC as well as with cerebellar regions that are part of DMN and fronto-parietal network (FPN) and with subcortical regions like the thalamus, hypothalamus, striatum (ventral and dorsal portions), and superior colliculus (**Fig. 1A**, **Supplementary Table 3**). Compared to HC subjects c-MCI patients showed no differences in the hippocampal FC with the rest if the brain (**Fig. 1B**). Finally, nc-MCI patients displayed extensive hyper-connectivity between the hippocampus and cerebellar areas that are functionally associated to DMN, FPN, and SN/CON (**Fig. 1C**, **Supplementary Table 4**).

**Figure 1.**
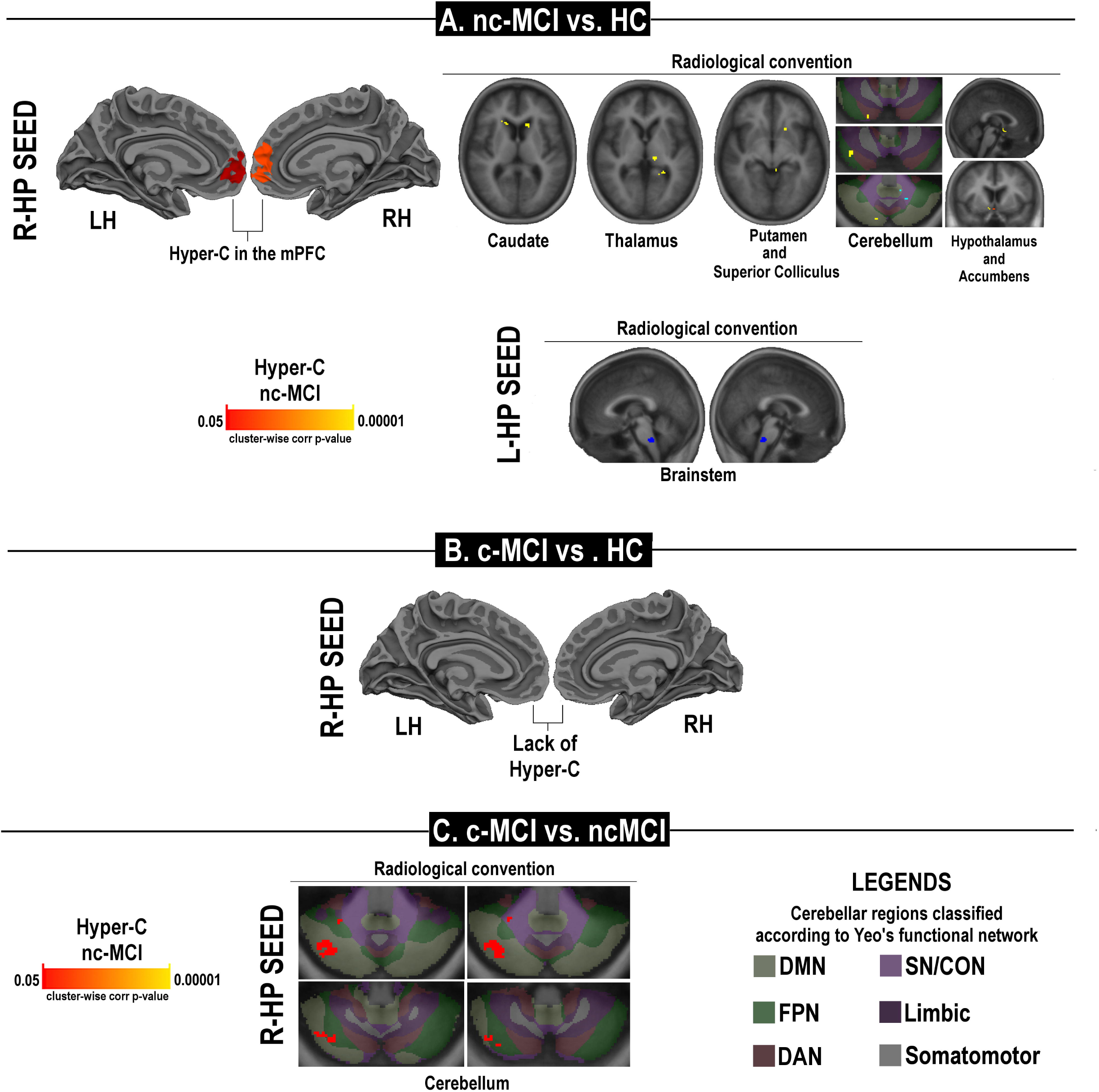
Statistical maps of differences in the hippocampal (HP) functional connectivity of MCI subsets. Panel A shows the comparison between nc-MCI and HC subjects; panels B and C show the comparison of c-MCI subjects with HC or nc-MCI subjects, respectively. The figure depicts areas with a cluster-wise probability below corrected p-value of 0.05. Pseudocolor scales indicate the statistical strength of the hyper-connectivity (Hyper-C) or hypo-connectivity (Hypo-C). Clusters changing from red to yellow or from dark blue to light blue are indicating either increased hyper-C or hypo-C, respectively. *Abbreviations:* DAN=Dorsal Attention Network; DMN=default-mode network; FPN=fronto-parietal network; SN/CON=Salience/Cingulo-Opercular Networks; L=left; LH=left hemisphere; mPFC=medial prefrontal cortex; R=right; RH=right hemisphere.

As far as the entorhinal cortex, nc-MCI patients showed hyper-connectivity in cerebellar areas that are part of the SN/CON, limbic and sensorimotor networks (**Fig. 2A**, **Supplementary Table 5**). Conversely, compared to HC (**Fig. 2B**, **Supplementary Table 6**) or nc-MCI subjects (**Fig. 2C**, **Supplementary Table 7**), c-MCI patients showed reduced connectivity with the lateral-occipital cortex, cortical and cerebellar regions that are part of attentive networks, brainstem, striatum, thalamus, and hypothalamus.

**Figure 2.**
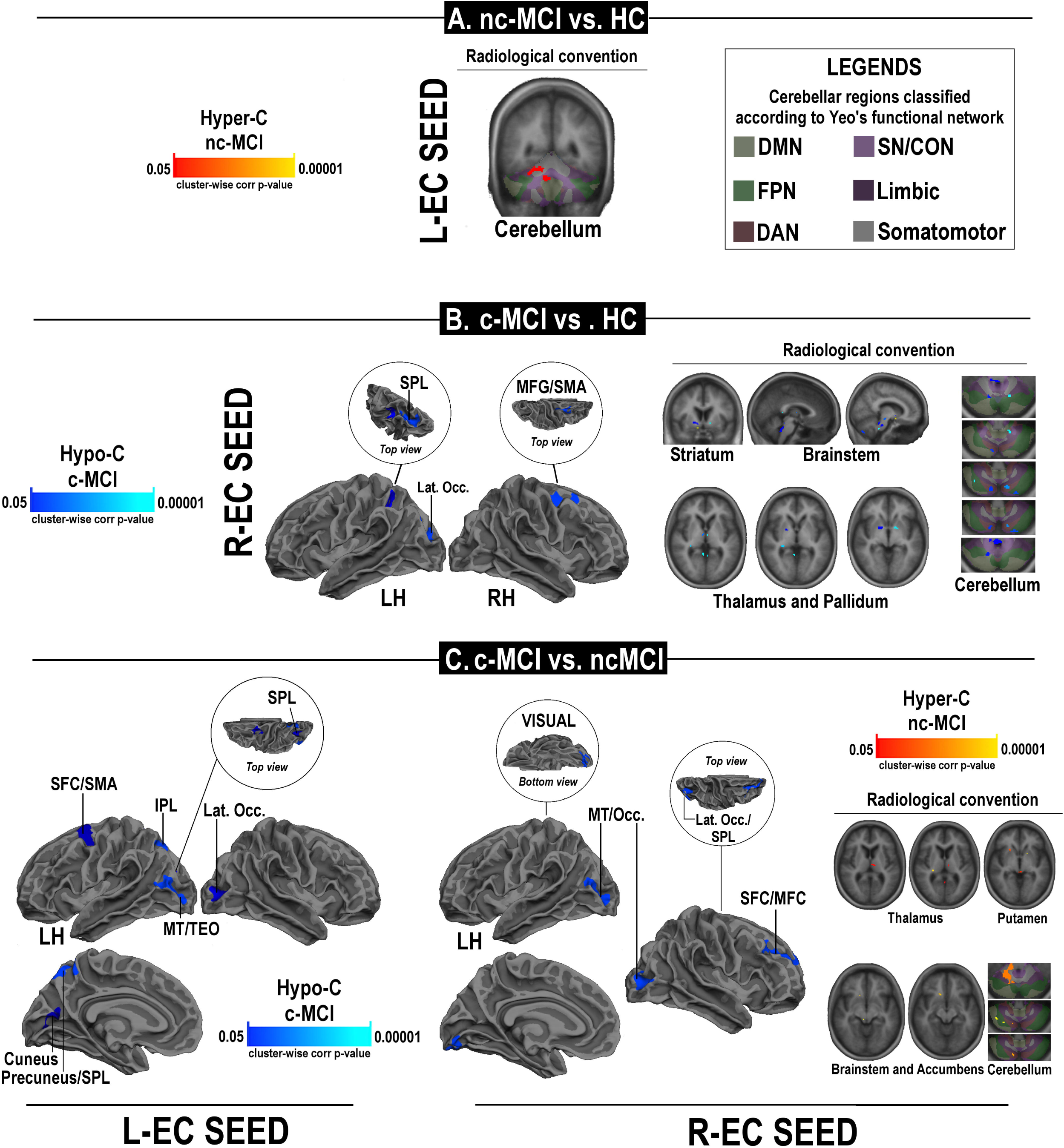
Statistical maps of differences in the entorhinal (EC) functional connectivity of MCI subsets. Panel A shows the comparison between nc-MCI and HC subjects; panels B and C show the comparison of c-MCI subjects with HC or nc-MCI subjects, respectively. The figure depicts areas with a cluster-wise probability below corrected p-value of 0.05. Clusters changing from red to yellow or from dark blue to light blue are indicating either increased hyper-C or hypo-C, respectively. *Abbreviations:* DAN=Dorsal Attention Network; DMN=default-mode network; FPN=fronto-parietal network; IPL= interior parietal lobe; L=left; LH=left hemisphere; MFG/SMA= middle frontal gyrus/supplementary motor area; mPFC=medial prefrontal cortex; R=right; RH=right hemisphere; SN/CON=Salience/Cingulo-Operacular Networks; SFC =superior frontal cortex; SPL=superior parietal lobe.

Compared to nc-MCI subjects, AD patients showed diffuse patterns of hippocampal hypo-connectivity with cortical and cerebellar regions that are mainly involved in the process belong to SN, DMN/PMN and sensorimotor network (**Fig. 3A**, **Supplementary Table 8**). Moreover, AD patients displayed entorhinal hypo-connectivity with brainstem, cortical regions of the DMN and SN/CON, and cerebellar areas that are part of the sensorimotor and attentional networks (**Fig. 4A**, **Supplementary Table 9**). In contrast, compared to c-MCI patients, AD individuals did not show differences in hippocampal FC but displayed entorhinal hypo-connectivity with the amygdala and brainstem (**Fig. 4B**, **Supplementary Table 10**).

**Figure 3.**
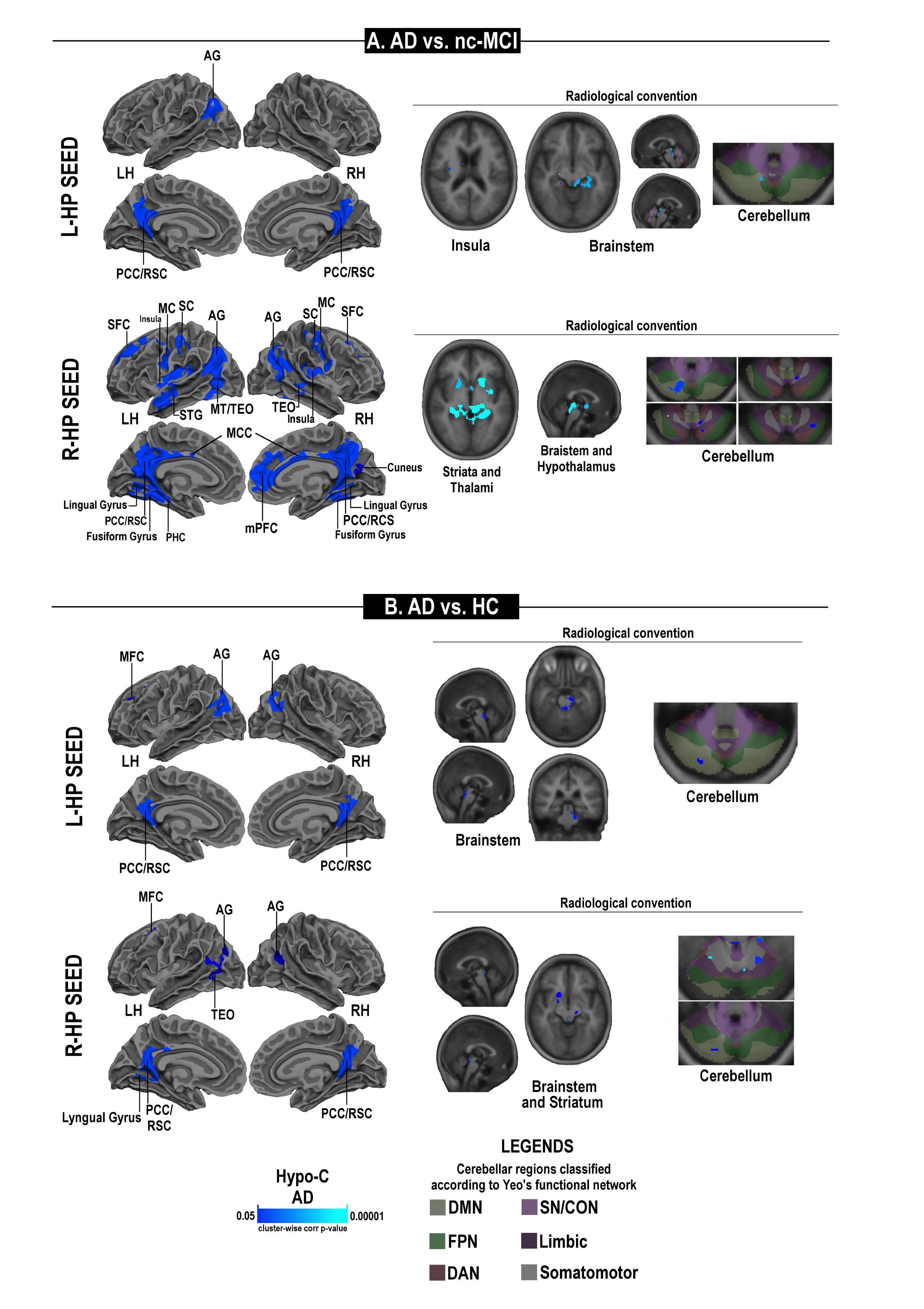
Statistical maps of differences in the hippocampal (HP) functional connectivity of AD patients. Significant hypo-connectivity is observed in AD patients when compared to nc-MCI (panel A) or HC subjects (panel B). The figure depicts areas with a cluster-wise probability below corrected p-value of 0.05. Pseudocolor scales, with clusters changing from dark blue to light blue, indicate the statistical strength of the hypo-connectivity (Hypo-C). *Abbreviations:* AG=angular gyrus; DAN=Dorsal Attention Network; DMN=default-mode network; FPN=fronto-parietal network; IPL= interior parietal lobe; L=left; LH=left hemisphere; MC=motor cortex; MCC=middle cingulate cortex; MFG/SMA= middle frontal gyrus/supplementary motor area; mPFC=medial prefrontal cortex; PCC/RSC=posterior cingulate cortex/retrosplenial cortex; PHC=parahippocampal cortex; R=right; RH=right hemisphere; SC=sensory cortex; SFC =superior frontal cortex; SN/CON=Salience/Cingulo-Operacular Networks; SPL=superior parietal lobe; STG=superior temporal gyrus.

**Figure 4.**
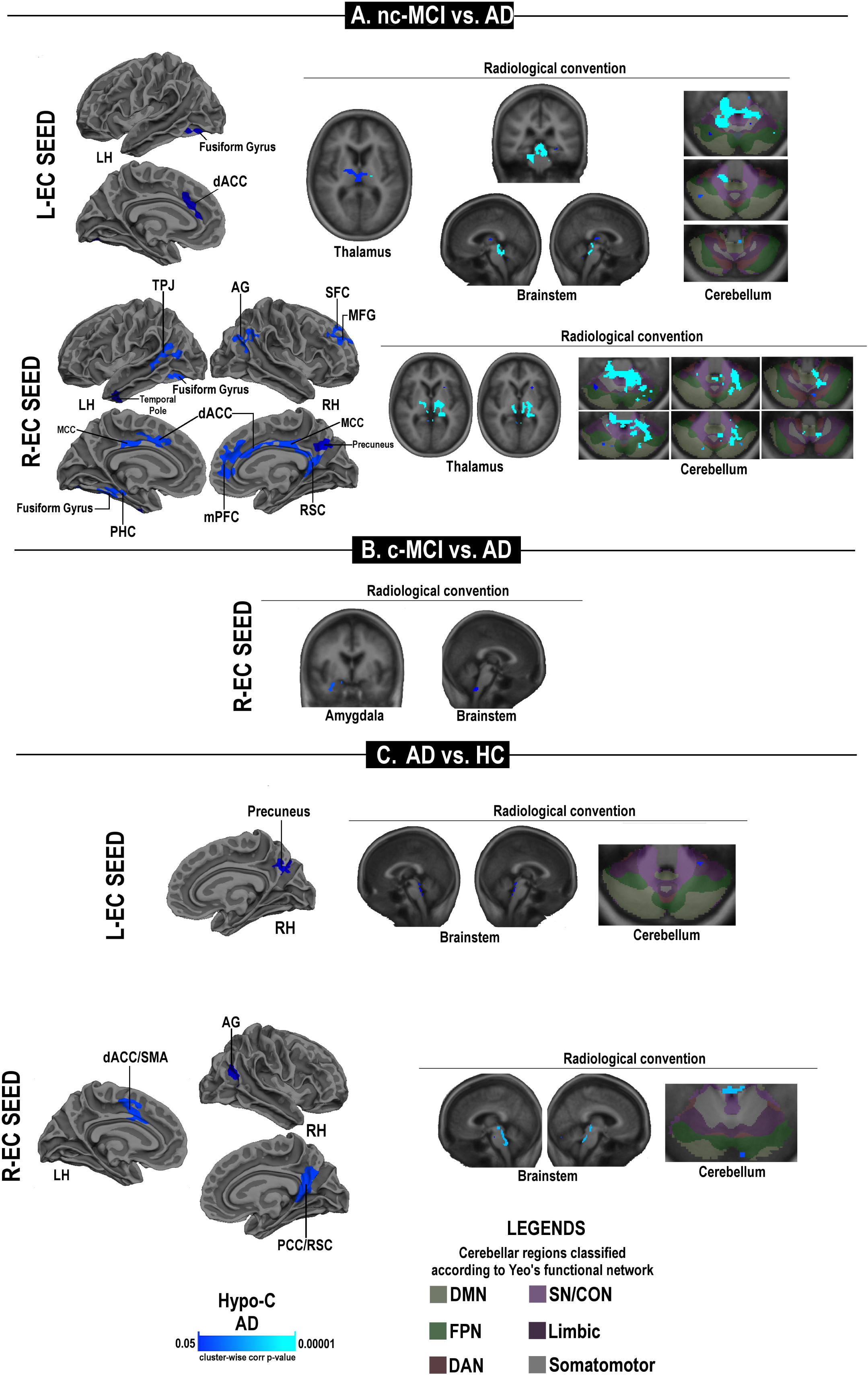
Statistical maps of differences in the entorhinal (EC) functional connectivity of AD patients. Significant hypo-connectivity is observed in AD patients when compared to nc-MCI (panel A), c-MCI (panel B), or HC subjects (panel C). The figure depicts areas with a cluster-wise probability below corrected p-value of 0.05. Pseudocolor scales, with clusters changing from dark blue to light blue, indicate the statistical strength of the hypo-connectivity (Hypo-C). *Abbreviations:* AG=angular gyrus; dACC=dorsal anterior cingulate cortex; DAN=Dorsal Attention Network; DMN=default-mode network; FPN=fronto-parietal network; IPL= interior parietal lobe; L=left; LH=left hemisphere; MC=motor cortex; MCC=middle cingulate cortex; MFC=middle frontal cortex; mPFC=medial prefrontal cortex; PCC/RSC=posterior cingulate cortex/retrosplenial cortex; PHC=parahippocampal cortex; R=right; RH=right hemisphere; SC=sensory cortex; SFC =superior frontal cortex; SMA/dACC=Supplementary Motor Area/dorsal Anterior Cingulate Cortex; SN/CON=Salience/Cingulo-Operacular Networks; SPL=superior parietal lobe; STG=superior temporal gyrus; TPJ=temporo-parietal junction.

### Relationship of rs-fMRI data with age, structures and CSF biomarkers of MCI patients

In the MCI group, by using a whole brain correlation analysis, we found that the features of hippocampal FC were associated with altered CSF levels of p-tau and amyloid while no relationships were found with age and alterations in the structural integrity of the hippocampi or cortices. Increased FC occurring between the hippocampus and DMN regions was negatively associated with levels of p-tau_181_ and p-tau_181_/Aβ_1–42_, and positively associated with Aβ_1–42_ levels (**Fig. 5**, **Supplementary Table 11**).

**Figure 5.**
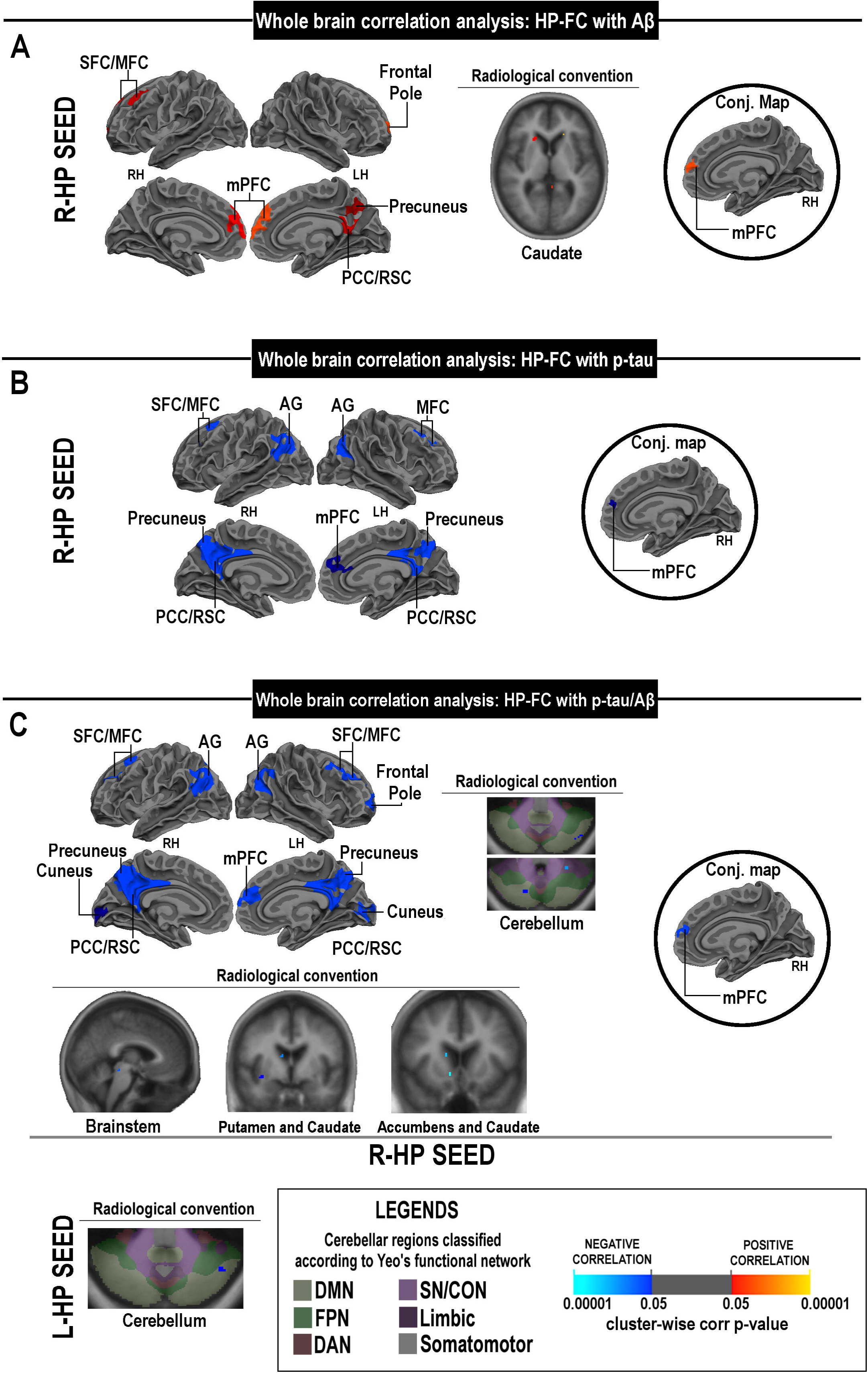
Whole-brain correlation analysis between the hippocampal (HP) functional connectivity and levels of CSF AD-related biomarkers. Maps depict clusters where the strength of the HP-FC significantly correlates with levels of Aβ_1–42_ (panel A), p-tau_181_ (panel B) and p-tau_181_/ Aβ_1–42_ (panel C). Black-boundaries circles highlight the results of the conjunction analyses. Each conjunction map shows the intersection of significant clusters that report the correlation between FC and CSF levels of biomarkers and clusters expressing significant FC differences in the comparisons between HC and MCI. Clusters changing from red to yellow or from dark blue to light blue are indicating either positive or negative correlations, respectively. *Abbreviations:* AG=angular gyrus; L=left; LH=left hemisphere; MC=motor cortex; MCC=middle cingulate cortex; MFC=middle frontal cortex; mPFC=medial prefrontal cortex; PCC/RSC=posterior cingulate cortex/retrosplenial cortex; R=right; RH=right hemisphere; SFC =superior frontal cortex.

The reduction of the strength of the right entorhinal FC was associated with cortical thinning of the right lingual gyrus and cuneus. No further associations were found between the entorhinal FC and another variable of interest (**Supplementary Fig. 6**).

## Discussion

In the present study, we investigated patterns of hippocampal and entorhinal FC in a cohort of HC, MCI and AD subjects. The rs-fMRI data were also evaluated in relation to the clinical progression of the study participants. Overall, the analysis indicates that AD patients showed a synergic and parallel process of hypo-connectivity localized in the hippocampus and entorhinal cortex. The analysis of the MCI subsets shows that, while the c-MCI patients are characterized by hypo-connectivity, nc-MCI patients exhibited hyper-connectivity of both entorhinal and hippocampal regions.

### Patterns of functional connectivity, cognitive status and structural damage in AD patients

Our AD subjects were characterized by the collapse of hippocampal and entorhinal connectivity, the decline in memory and executive skills, and the presence of marked signs of cortical and subcortical atrophy. These findings confirm the notion that macro-structural damage severely impairs global communication efficiency, prevents the adaptive functional reorganization of the brain networks, and ultimately sets the stage for the disease progression (Hillary and Grafman 2017). Of note, we observed a hypo-connectivity in the angular gyrus and retrosplenial/posterior cingulate cortex, two areas that are strictly involved in memory retrieval and intimately connected to the hippocampus and entorhinal cortex (Eichenbaum 2017; Sestieri et al. 2017). It is therefore possible that, in AD subjects, the reduced connectivity of the hippocampus and entorhinal cortex represents the functional correlate of the defective episodic memory retrieval that is typically found in the disease.

### Patterns of functional connectivity, cognitive status and structural features in nc-MCI patients

Our study shows that nc-MCI patients exhibited hippocampal and entorhinal hyper-connectivity, relative preservation of cognitive functions and brain structures, and non-pathological levels of the AD-related CSF biomarkers. The findings are in line with previous studies showing patterns of increased hippocampal FC occurring in healthy, but at-risk for AD, individuals as well as in MCI patients (Bookheimer et al. 2000; Bondi et al. 2005; Hamalainen et al. 2007; Kircher et al. 2007; Das et al. 2013; Putcha et al. 2011). The hippocampal hyperactivity exhibited by MCI patients is controversial in value. While some authors have proposed that the process plays a compensatory role and helps to maintain cognitive performances (Sperling et al. 2009; Mormino et al. 2012; Oh and Jagust, 2013; Huijbers et al. 2015), others have considered the phenomenon disadvantageous and set to promote cognitive impairment (Das et al. 2013; Pasquini et al. 2015).

Our results that indicate the presence of enhanced FC between the hippocampus, thalamus, striatum, and mPFC lend support to the “compensatory hypothesis”. The thalamus is a structural and functional hub of the communication occurring between the hippocampus and mPFC, thereby supporting strategic cognitive functions, including memory consolidation (Ferraris et al. 2018) (Eichenbaum 2017). The striatum, along with the PFC, is also implicated in the modulation of memory retrieval (Scimeca JM and D Badre 2012). The mPFC is part of an integrated system (DMN) that sustains the global communication and meta-stability of the brain (Hellyer et al. 2014) and, ultimately, modulates a wide-range of high-order cognitive functions as well as the resilience against neurodegenerative processes (Hillary and Grafman 2017). The hyper-connectivity with the mPFC, is in line with different modelizations of brain ageing-related dynamics (i.e., HERA, HAROLD, PASA, CRUNCH, STAC, GOLDEN Aging) that postulate an increased engagement of the prefrontal brain regions to compensate for the functional decline of the posterior regions (Tulving et al. 1994; Cabeza et al. 1997; Davis et al. 2008) (Schneider-Garces et al. 2010; Park and Reuter-Lorenz 2009; Reuter-Lorenz and Park 2014; Fabiani M 2012). The compensatory hypothesis fits with evidence indicating that the mPFC and the hippocampus, the two areas where we observed increased FC, are tightly interconnected by bidirectional projections that are structurally and functionally integrated. The oscillatory synchronic activity between these two regions supports the organization and processing of the episodic memory (Eichenbaum 2017). The mPFC is strategic for memory as the area receives information on contextual cues from the anterior hippocampus and, in turn, indirectly sends the information, via thalamus and perirhinal/entorhinal cortices, to the posterior hippocampus (**Fig. 6**). In this context, the hippocampus acts as a key region set to control the memory organization and encoding, whereas the mPFC is implicated in the retrieval of context-appropriate memory engrams, the suppression of distractors or interference and the switching or selection of episodic memories according to contextual rules (Eichenbaum 2017). Furthermore, the presence of altered connectivity between the mPFC and the hippocampus impairs the object-place and temporal-order memory and leads to severe impairment in conditional visual discrimination as well as to learning and memory deficits related to defective suppression of irrelevant memory engrams (Eichenbaum 2017; Barker et al. 2007).

**Figure 6.**
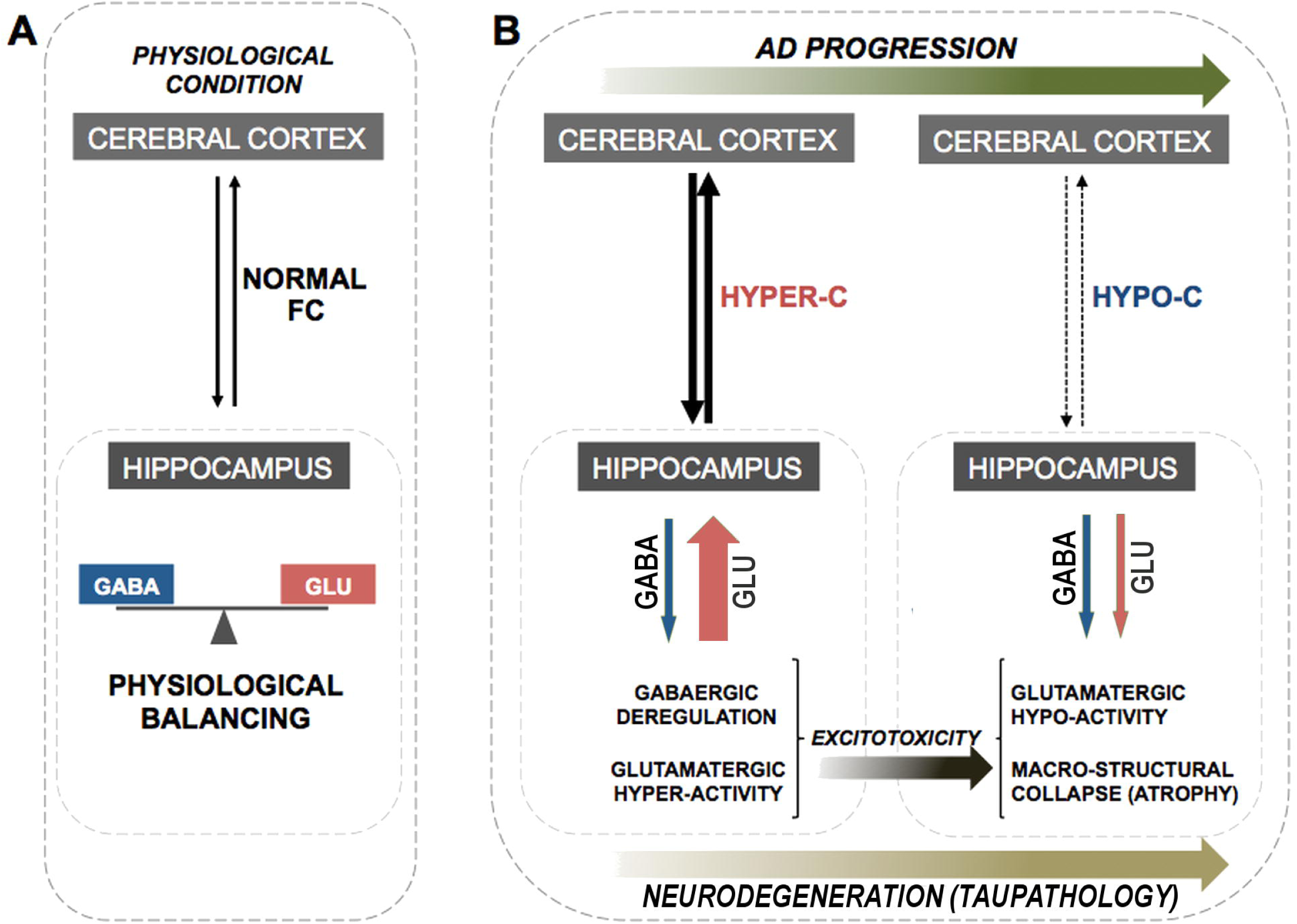
Diagram of the neural system underlying episodic memory. The superficial layers II-III of the entorhinal cortex (EC) carry unimodal/multimodal cortical information from the cortical associative areas (via perirhinal cortex, PRC or parahippocampal cortex, PHC) to the hippocampus (HP). The deep layers V-VI of the lateral entorhinal cortex (LEC), through the cingulum bundle, project back signals from the posterior hippocampus (pHP) to the PHC and the posterior regions of default-mode network (DMN). These pathways are promoting the integration of spatial information and the representation of retrieved events, respectively. The anterior hippocampus (aHP) carries global information on contextual cues to the mPFC. The mPFC then selects memory engrams within the hippocampus, manages the rule-guided switching between memory-related strategies and supervises the memory retrieval process by exerting a top-down (cognitive or strategic) modulation of the activity of the posterior hippocampus through the LEC and PRC pathways. Further abbreviations: ACC/mPFC=anterior cingulate cortex/mPFC; AG=angular gyrus; MEC=medial entorhinal cortex; OFC=orbitofrontal cortex; PCC=posterior cingulate cortex; PCN=precuneus; RSC=restrosplenial cortex; TE/TEO=temporoccipital association areas V4=visual cortex.

Interestingly, nc-MCI individuals were characterized by hippocampal hyper-connectivity with the hypothalamus, superior colliculus, and cerebellar areas that are functionally associated to the DMN and the FPN/DAN. These subjects also displayed hyper-connectivity between the entorhinal cortex and cerebellar regions that are part of the somatosensory network. Thus, our findings support a close functional connection between the hippocampus and cerebellum. The interplay between these regions, through circuits that involve the entorhinal cortex, hypothalamus, superior colliculus, and thalamus (including the cerebello-thalamo-cortical and cortico-ponto-cerebellar pathways), is strategic for the modulation of cognitively relevant prefrontal and parietal activities (Yu W and E Krook-Magnuson 2015). Moreover, the involvement of the cerebellum is intriguing as recent evidence indicates that the region acts as a critical hub for the control of a wide range of cognitive processes encompassing language, visual-spatial, executive, and working memory processes (Stoodley 2012). Thus, the hyper-connectivity between the hippocampus and cerebellum is functionally relevant and potentially associated with the relative preservation of the high-order cognitive functions of nc-MCI individuals.

Thus, from a theoretical standpoint, the increased FC that we observed in nc-MCI subjects may help to transiently cope with, and counteract, the undergoing neurodegenerative process and related cognitive impairment.

### Patterns of functional connectivity, cognitive status and structural alterations in c-MCI patients

In contrast to nc-MCI subjects, c-MCI patients did not show signs of hippocampal hyper-connectivity with mPFC. This lack of hyper-connectivity may result in the reduced compensatory engagement of prefrontal areas and a more severe cognitive decline. In line with previous MRI studies (Grundman et al. 2002; Jack CR Jr. et al. 2004; Apostolova et al. 2006; Henneman et al. 2009), c-MCI patients showed hippocampal atrophy. These patients also showed entorhinal hypo-connectivity with cortical and cerebellar regions that take part in the modulation of long-term memory and attentional systems. Entorhinal hypo-connectivity, in particular, should be considered in relation to the role played by the superficial layers, II-III, of this region (**Fig. 6**). These layers act, in fact, as relay stations that carry, through the perirhinal or parahippocampal cortices, unimodal/multimodal cortical information from cortical associative areas to the hippocampus (Canto et al. 2008; Ranganath and Ritchey 2012). Moreover, the deep layers, V-VI, of the lateral entorhinal cortex send projections from the posterior hippocampus, via the cingulum, to the parahippocampus and the cortical areas involved in attentional networks and the DMN/PMN bundle (Kahn et al. 2008; Lacy and Stark 2012; Libby et al. 2012). These circuits promote the integration of spatial information as well as the representation of retrieved events (Preston and Eichenbaum 2013; Vann et al. 2009). It is therefore conceivable that the c-MCI reduced FC within the DMN/PMN and attentional/associative networks represents a functional marker of underlying alterations that occur before the onset and development of AD.

Overall, these data are in agreement with neuropathological evidence indicating that the entorhinal cortex and the hippocampus are the first brain regions to display tau-pathology and neurodegeneration in the course of AD (Braak et al. 1994). In line with this notion, our c-MCI patients showed decreased levels of Aβ_1–42_ and increased levels of t-tau, p-tau_181_, t-tau/Aβ_1–42_, and p-tau_181_/Aβ_1–42_, CSF alterations that went along with the presence of more profound memory deficits.

The correlation between altered CSF features and mPFC-related modifications is in line with studies showing that the decreased FC between central hubs of the DMN correlates with enhanced Aβ deposition (Buckner RL et al. 2005) (Elman et al. 2016; Foster et al. 2018; Grothe et al. 2016; Koch et al. 2010; Mutlu et al. 2017; Mormino et al. 2011; Palmqvist et al. 2017). The link between the presence of hyper-connectivity and enhanced signs of tau-pathology is less explored. However, fMRI/PET studies have recently shown that increased levels of tau-related pathology lead to a progressive decline of the brain FC (Cope et al. 2018; Hoenig et al. 2018; Jones et al. 2016; Schultz et al. 2017; Sepulcre et al. 2017) and the activation of the DMN in particular (Hoenig et al. 2018). These findings go along with evidence indicating that the loss of hippocampal GABAergic inter-neurons is closely associated with the appearance of enhanced signs of tau-pathology (Levenga et al. 2013). These processes may accelerate an ongoing pattern of hippocampal hyperactivity, micro/ macro-structural damage, atrophy, and the progression of cognitive and behavioral disorders (Gilani et al. 2014; Jones et al. 2016; Schmitz et al. 2017; Schobel et al. 2013).

We, therefore, propose a “work in progress” model by which a pattern of altered hippocampal FC may identify the degenerative processes that are driving MCI subjects to become AD patients. It is conceivable that the functional/dysfunctional value of the process varies upon different stages of the AD-related spectrum. At the MCI stage, the down-regulation of GABAergic neurotransmission may unleash a glutamatergic overdrive that promotes, a transiently beneficial, enhancement of the hippocampal activity that improves the functional coupling of the region with the cortex. This up-regulation is, for a while, advantageous and leads to increased communication between the hippocampus and cortical areas, like the mPFC, that are critically involved in the brain meta-stability and protection of cognitive functioning and, when deprived of activity, became more susceptible to AD and amyloidosis. This hypothesis is supported by the preservation of cognitive functions that we find in our nc-MCI patients. In the long run, the hippocampal increased FC may, however, set the stage for an enhanced, activity-dependent, damage of the region that is likely to be carried out by a combination of increased tau-related pathology [as we indirectly observed in the CSF of our c-MCI patients (**Table 2**)] and excitotoxicity, two processes ultimately leading to decreased FC and clinical conversion to AD (**Fig. 7**).

**Figure 7.**
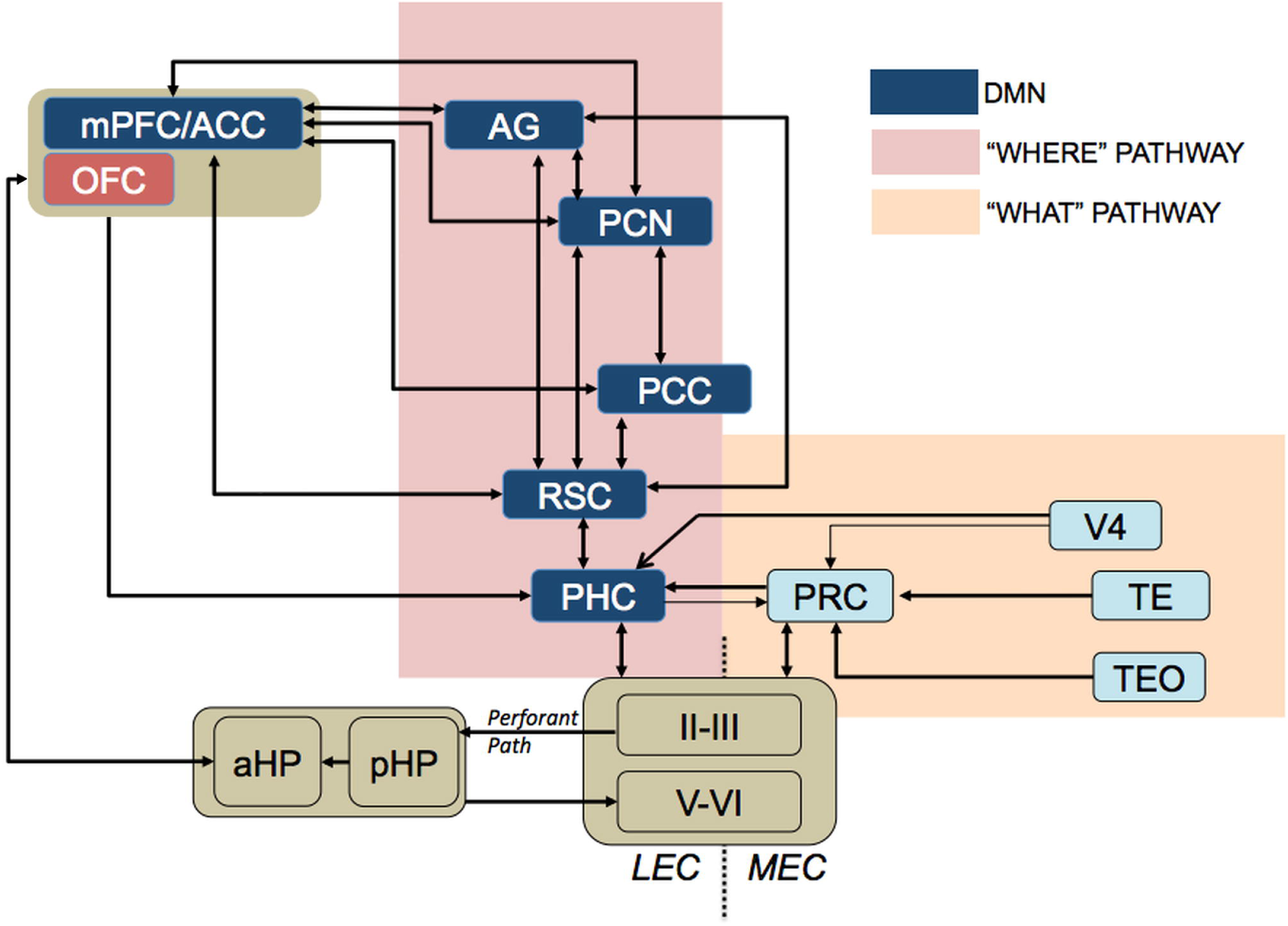
Model of the bi-modal hippocampal connectivity that occurs at different stages of Alzheimer’s Disease spectrum. Upon physiological conditions, the functional connectivity (FC) of the hippocampus results from a balanced and synergistic activation of GABA-mediated inhibition and glutamate-mediated excitation (Panel A). At the MCI stage, the reduced and pathology-driven, activation of GABAergic interneurons unleashes a glutamatergic overdrive that sets the stage for a transient compensatory enhancement of hippocampal activity promoting improved functional coupling with the cortex (Panel B). However, in the long run, this increased glutamate-dependent FC may trigger an enhanced process of activity-dependent hippocampal damage. The damaging process is driven by the increased accumulation of tau-pathology and excitotoxicity, ultimately, favoring the development of macro/micro-structural damage, functional collapse of the region, increased cognitive deficits, and the progression to AD. Abbreviations: Hyper-C= hyper-connectivity; Hypo-C= hypo-connectivity; GLU=glutamate.

In conclusion, the study identifies the functional correlates of alterations that occur in patients at the early stages of the AD-related spectrum. Our study has some limitations. For instance, it should be underlined that all the study subjects are highly educated individuals who likely possess significant levels of cognitive reserve. Furthermore, the neuropsychological tests employed in the ADNI database are skewed toward the investigation of mnemonic domains and do not allow a detailed analysis of visuospatial and attentional domains. Our working hypothesis warrants future longitudinal investigation. For instance, the model should be tested and further validated by investigating changes in functional and structural connectivity in relation to ongoing processes of amyloid and tau deposition as assessed by PET imaging.

## Acknowledgments

This work was supported by research grants from the Italian Department of Education (PRIN 2011; 2010M2JARJ_005) and the Italian Department of Health (RF-2013–02358785 and NET-2011-02346784-1).

Data collection and sharing for this project was funded by the ADNI (National Institutes of Health Grant U01 AG024904). ADNI is funded by the National Institute on Aging, the National Institute of Biomedical Imaging and Bioengineering, and through generous contributions from the following: AbbVie, Alzheimer’s Association; Alzheimer’s Drug Discovery Foundation; Araclon Biotech; BioClinica, Inc.; Biogen; Bristol-Myers Squibb Company; CereSpir, Inc.; Cogstate; Eisai Inc.; Elan Pharmaceuticals, Inc.; Eli Lilly and Company; EuroImmun; F. Hoffmann-La Roche Ltd and its affiliated company Genentech, Inc.; Fujirebio; GE Healthcare; IXICO Ltd.;Janssen Alzheimer Immunotherapy Research & Development, LLC.; Johnson & Johnson Pharmaceutical Research & Development LLC.; Lumosity; Lundbeck; Merck & Co., Inc.;Meso Scale Diagnostics, LLC.; NeuroRx Research; Neurotrack Technologies; Novartis Pharmaceuticals Corporation; Pfizer Inc.; Piramal Imaging; Servier; Takeda Pharmaceutical Company; and Transition Therapeutics. The Canadian Institutes of Health Research is providing funds to support ADNI clinical sites in Canada. Private sector contributions are facilitated by the Foundation for the National Institutes of Health (www.fnih.org). The grantee organization is the Northern California Institute for Research and Education, and the study is coordinated by the Alzheimer’s Therapeutic Research Institute at the University of Southern California. ADNI data are disseminated by the Laboratory for Neuro Imaging at the University of Southern California.

The authors are grateful to all the members of the Molecular Neurology Unit for helpful discussions and to Dr. Domenico Ciavardelli for helping with the statistical analysis.

---

**Supplementary Figure 1.**
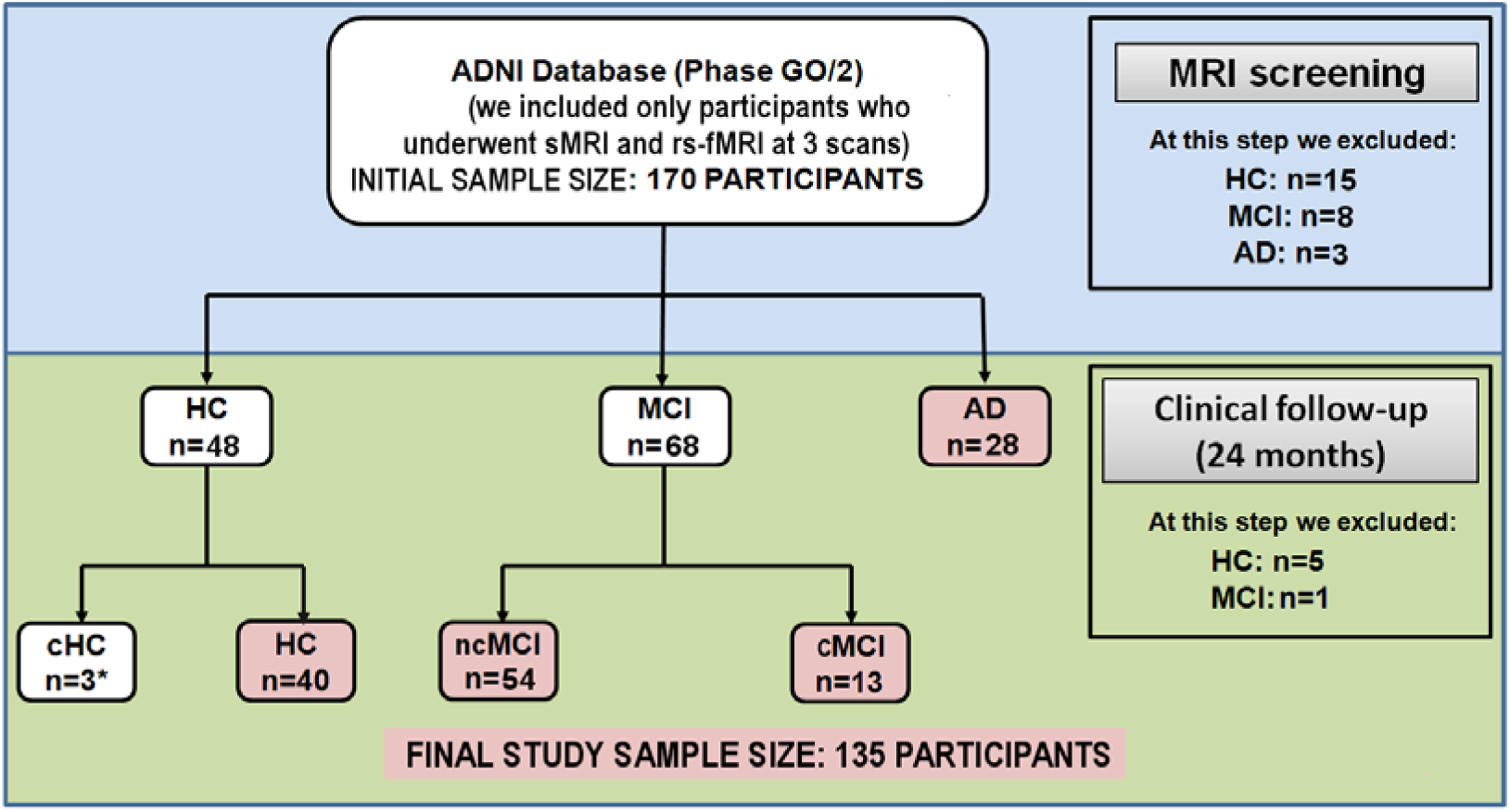
Flow-chart of the study sample selection. Abbreviations: HC=healthy control subjects; MCI=Mild Cognitive Impairment; AD=Alzheimer’s Disease; c=converters; nc=patients who did not convert to AD.

**Supplementary Figure 2.**
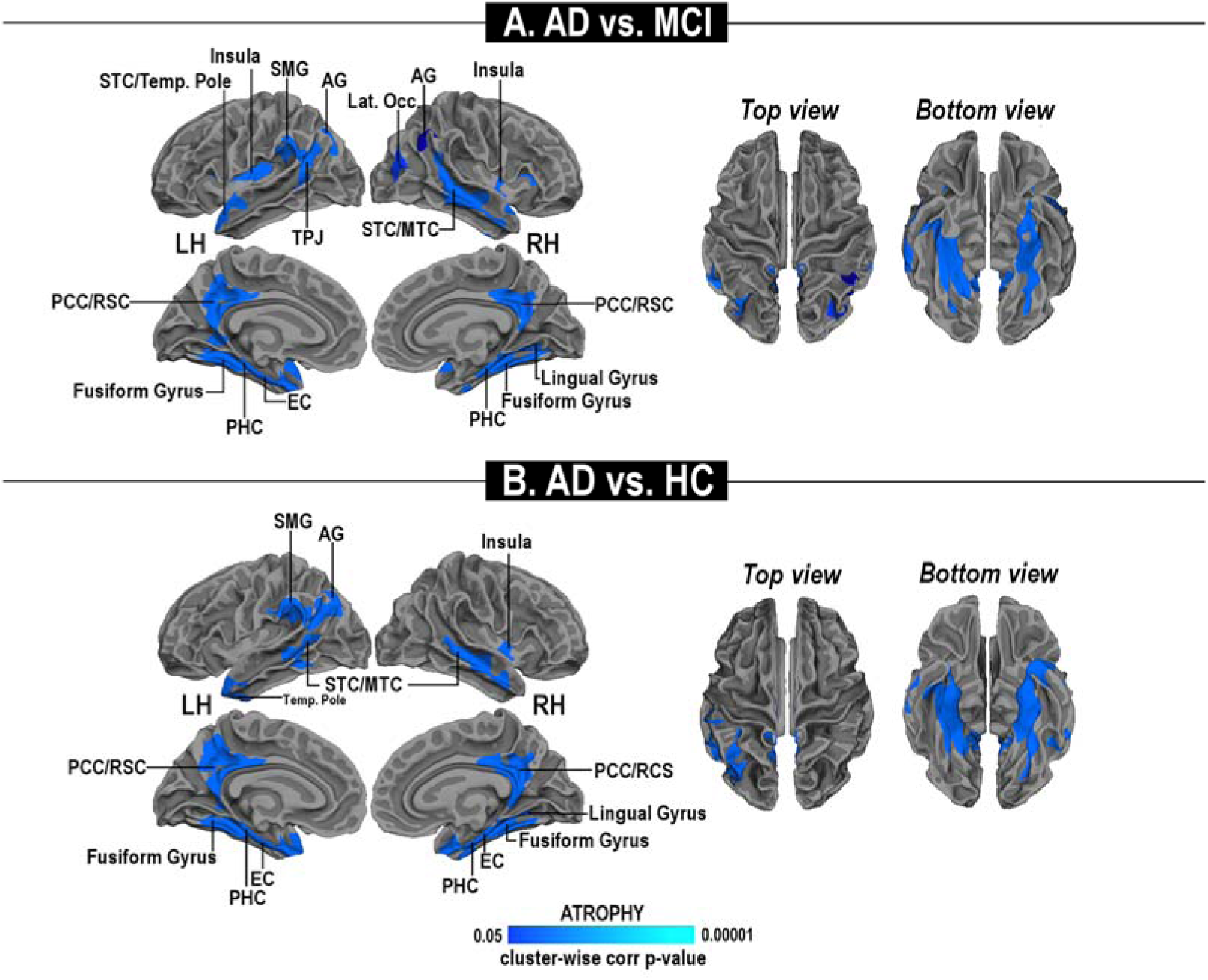
Cortical thickness analysis of the study groups (model I). Statistical maps depict differences in cortical thickness of Alzheimer’s Disease (AD), Mild Cognitive Impairment (MCI) or Healthy elderly (HC) subjects. Significant cortical thinning is observed in AD patients when compared to HC (panel A) or MCI subjects (panel B). No statistically significant differences are found in the HC vs. MCI comparison. The figure depicts areas with a cluster-wise probability below corrected p-value of 0.05. Pseudocolor scales indicate the statistical value of the regional cortical thinning (light blue =low and dark blue=high). LH=left hemisphere; RH=right hemisphere; L=left; R=right.

**Supplementary Figure 3.**
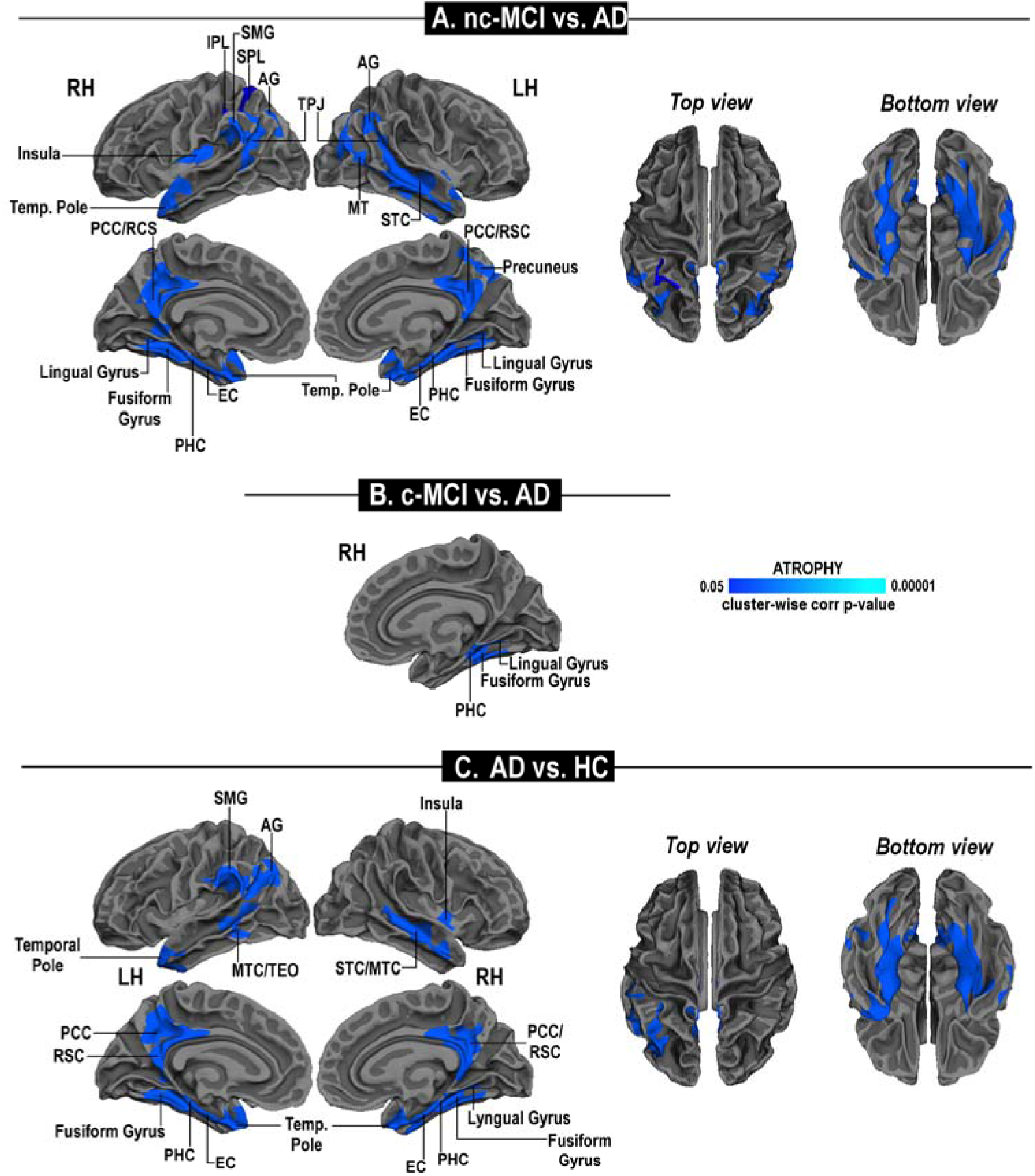
Cortical thickness analysis including the MCI subtypes (model II). Statistical maps depict differences in cortical thickness of AD and MCI subsets. Panel A shows the comparison between nc-MCI and c-MCI patients. Panel B and C show, respectively, the cortical thinning in AD patients when compared to nc-MCI or c-MCI subjects. The figure depicts areas with a cluster-wise probability below corrected p-value of 0.05. Pseudocolor scales indicate the statistical value of the regional cortical thinning (light blue =low and dark blue=high). LH=left hemisphere; RH=right hemisphere; L=left; R=right.

**Supplementary Figure 4.**
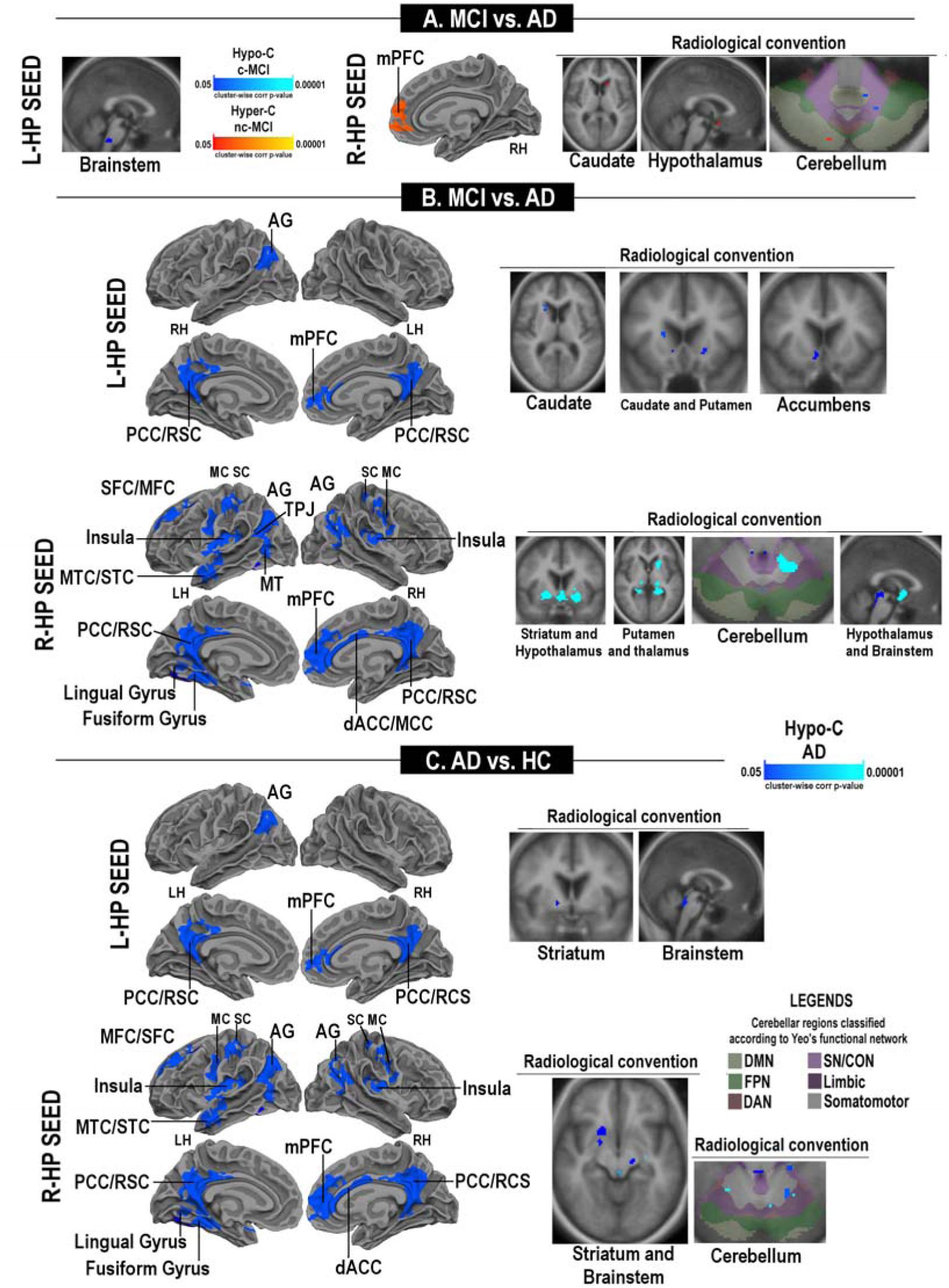
Statistical maps of differences in the hippocampal (HP) functional connectivity of AD patients (model I). Panel A shows the comparison between HC and MCI patients; panels B and C show hypo-connectivity in AD patients when compared to MCI or HC subjects, respectively. The figure depicts areas with a cluster-wise probability below corrected p-value of 0.05. Clusters changing from red to yellow or from dark blue to light blue are indicating either increased hyper-C or hypo-C, respectively. RH=right hemisphere; L=left; R=right.

**Supplementary Figure 5.**
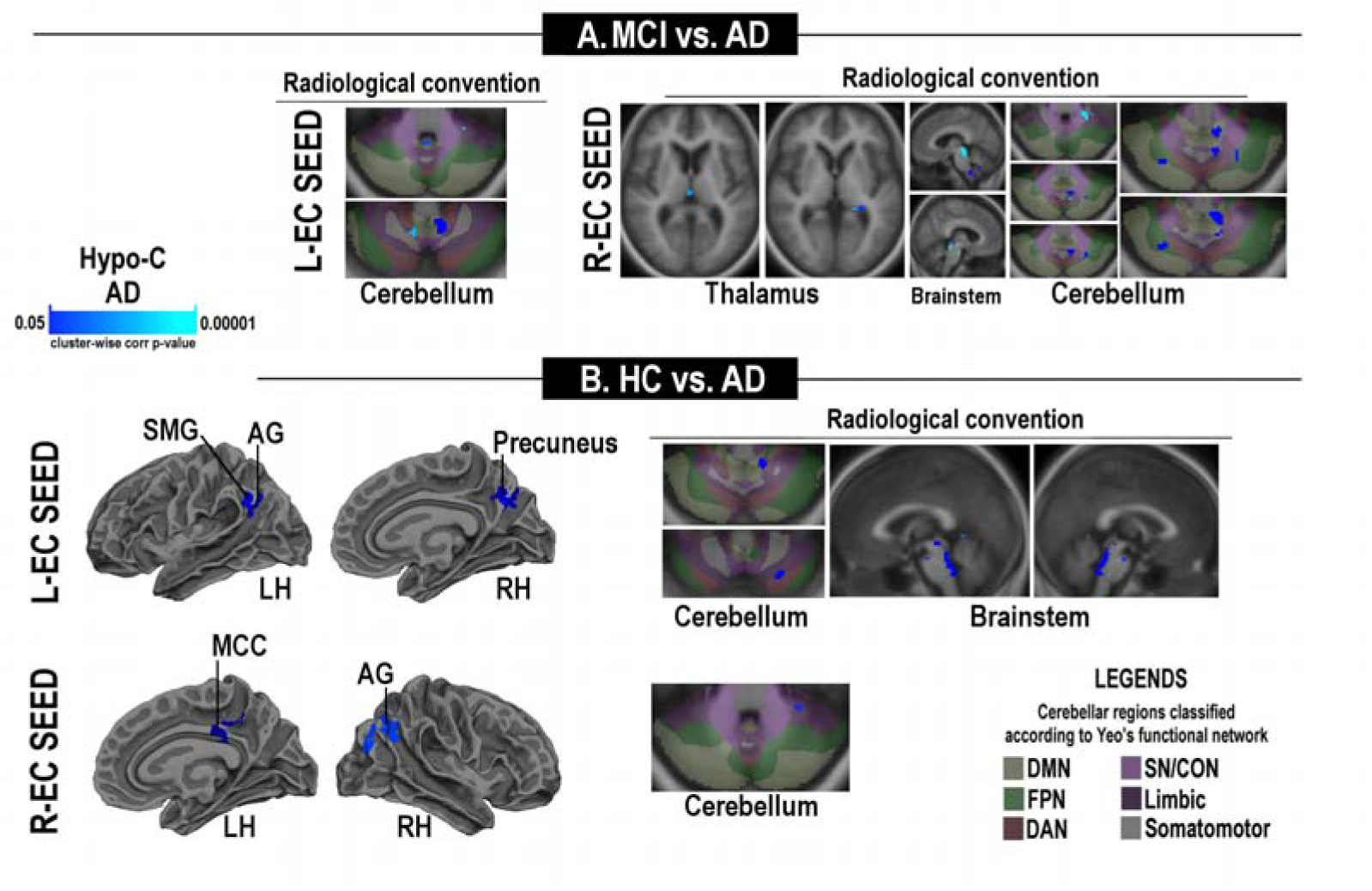
Statistical maps of differences in the entorhinal functional connectivity in AD patients (model I). Panels A and B show hypo-connectivity in AD patients when compared to MCI or HC subjects, respectively. The figure depicts areas with a cluster-wise probability below corrected p-value of 0.05. Pseudocolor scale, with clusters changing from dark blue to light blue, indicates the statistical strength of the hypo-connectivity (Hypo-C). LH=left hemisphere; RH=right hemisphere; L=left; R=right.

**Supplementary Figure 6.**
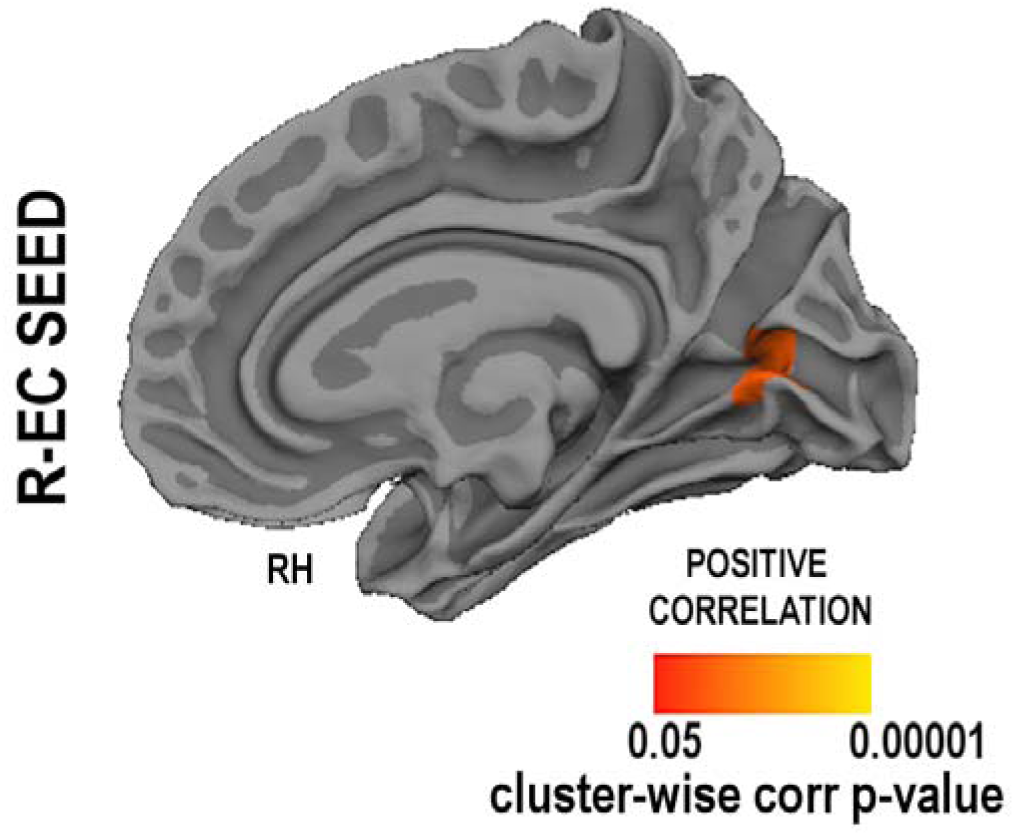
Correlation maps of entorhinal functional connectivity (FC) with cortical thickness in MCI patients. The figure depicts the clusters where FC differences were significantly correlated with levels of p-tau181 (panel A), the p-tau181/A_β1–42_ ratio (panel B). No further association was found. Clusters changing from dark blue to light blue are indicating negative correlations. LH=left hemisphere; RH=right hemisphere; L=left; R=right.

**Supplementary Table 1.** Hippocampal (HP) functional connectivity analysis: comparison between AD patients and HC subjects (II model).

| Cortical analysis |  |  |  |  |  |  |  |  |  |
| --- | --- | --- | --- | --- | --- | --- | --- | --- | --- |
| Regions included in the cluster | Size (mm<sup>2</sup>) | MNI - X | MNI - Y | MNI - Z | CWP | CWP Low | CWP Hi | NVtxs | WghtVtx |
| Seed: R-HP |  |  |  |  |  |  |  |  |  |
| R-PCC/RSC | 1209.3 | 11.4 | -46.2 | 3 | 0.0003 | 0 | 0.0006 | 2960 | 8479.73 |
| R-AG | 532.51 | 41.9 | -63.4 | 23.9 | 0.00389 | 0.0027 | 0.00539 | 1038 | 2881.34 |
| L-TEO | 403.89 | 51 | -37.1 | 7 | 0.03586 | 0.03176 | 0.03996 | 955 | 2409.41 |
| L- PCC/RSC | 1478.45 | -7.4 | -50.5 | 8.7 | 0.0003 | 0 | 0.0006 | 3485 | 10253.9 |
| L-MFC | 617.67 | -28.2 | 9.6 | 50.1 | 0.0006 | 0 | 0.0012 | 1128 | 3522.9 |
| L-Lat. Occ. Cort. | 586.34 | -40.5 | -73.5 | 19.7 | 0.0015 | 0.0006 | 0.0024 | 1124 | 2685 |
| L-AG | 517.16 | -35.1 | -79.8 | 31.5 | 0.00629 | 0.00449 | 0.00808 | 859 | 2537.95 |
| Seed: L-HP |  |  |  |  |  |  |  |  |  |
| R-PCC/RSC | 1106.35 | 5.3 | -65.9 | 29 | 0.0003 | 0 | 0.0006 | 2664 | 7827.46 |
| R-AG | 731.26 | 47.4 | -62 | 23.2 | 0.0003 | 0 | 0.0006 | 1228 | 3397.07 |
| L-AG | 1476.18 | -37 | -77.1 | 31.3 | 0.0003 | 0 | 0.0006 | 2594 | 7957.31 |
| L-PCC/RSC | 1023.32 | -5.5 | -58.6 | 19.1 | 0.0003 | 0 | 0.0006 | 2422 | 7271.15 |
| L-MFC | 582.76 | -34.9 | 13.4 | 52.3 | 0.0015 | 0.0006 | 0.0024 | 1104 | 2756.86 |
| Subcortical analysis |  |  |  |  |  |  |  |  |  |
| Regions included in the cluster | Size (mm<sup>2</sup>) | MNI - X | MNI - Y | MNI - Z | GRF CWP | Cortical network associated to cerebellar region |  |  |  |
| Seed: R-HP |  |  |  |  |  |  |  |  |  |
| R-Ventral DC | 16 | 18 | -21 | -13 | 0 |  |  |  |  |
| L-Cerebellum | 80 | -22 | -53 | -21 | 0.0062412 | SMN |  |  |  |
| L-Brainstem | 96 | 0 | -37 | -19 | 0.0112608 |  |  |  |  |
| R-Putamen | 1184 | 16 | 13 | -11 | 0.0309982 |  |  |  |  |
| L-Hippocampus | 1368 | -22 | -39 | -1 | 0.0383811 |  |  |  |  |
| R-Cerebellum | 104 | 24 | -79 | -29 | 0.0424053 | DMN |  |  |  |
| Seed: L-HP |  |  |  |  |  |  |  |  |  |
| Brainstem | 2080 | -10 | -21 | -25 | 0.0110809 |  |  |  |  |
| R-Striatum | 64 | 16 | 3 | -5 | 0.0287001 |  |  |  |  |
| R-Cerebellum | 104 | 26 | -79 | -35 | 0.04354 | DMN |  |  |  |

*Abbreviations:* AG=angular gyrus; CWP=cluster wise p-value; CWP Low and CWP Hi=90% confidence interval for CWP; NVtxs: number of vertices in cluster; DC= ventral diencephalon; DMN=Default-Mode Network; EC=entorhinal cortex; R=right; PCC/RSC=posterior cingulate cortex/retrosplenial cortex; SMN=Sensorimotor Network.

**Supplementary Table 2.** Entorhinal (EC) functional connectivity analysis: comparison between AD patients and HC subjects (II model).

| Cortical analysis |  |  |  |  |  |  |  |  |  |
| --- | --- | --- | --- | --- | --- | --- | --- | --- | --- |
| Regions included in the cluster | Size (mm<sup>2</sup>) | MNI - X | MNI - Y | MNI - Z | CWP | CWP Low | CWP Hi | NVtxs | WghtVtx |
| Seed: R-EC |  |  |  |  |  |  |  |  |  |
| R-AG | 1013.49 | 47.5 | -58.4 | 38.2 | 0.0003 | 0 | 0.0006 | 1823 | 1013.49 |
| R-PCC/RSC | 469.1 | 4.5 | -28.1 | 31.8 | 0.003 | 0.0018 | 0.00419 | 1161 | 469.1 |
| R-Precuneus | 435.09 | 5.3 | -66.8 | 31.1 | 0.00838 | 0.00629 | 0.01046 | 804 | 435.09 |
| L-dACC/SMA | 680.17 | -9.5 | 9.8 | 35.2 | 0.0003 | 0 | 0.0006 | 1632 | 4724.78 |
| Seed: L-EC |  |  |  |  |  |  |  |  |  |
| R-Precuneus | 487.23 | 5.3 | -66.8 | 31.1 | 0.0027 | 0.0015 | 0.00389 | 999 | 2658.04 |
| Sub-cortical analysis |  |  |  |  |  |  |  |  |  |
| Regions included in the cluster | Size (mm<sup>2</sup>) | MNI - X | MNI - Y | MNI - Z | GRF CWP | Cortical network associated to cerebellar region |  |  |  |
| Seed: R-EC |  |  |  |  |  |  |  |  |  |
| Brainstem | 4232 | 2 | -43 | -37 | 0.0001012 |  |  |  |  |
| R-Cerebellum | 80 | -10 | -87 | -23 | 0.0291045 | DMN |  |  |  |
| Seed: L-EC |  |  |  |  |  |  |  |  |  |
| L-Cerebellum | 72 | -32 | -39 | -33 | 0.0071834 | Limbic Network |  |  |  |
| Brainstem | 104 | 0 | -35 | -15 | 0.0114115 |  |  |  |  |
| Brainstem | 152 | 4 | -43 | -39 | 0.0168259 |  |  |  |  |
| Brainstem | 176 | -4 | -41 | -29 | 0.029145 |  |  |  |  |

*Abbreviations:* AG=angular gyrus; CWP=cluster wise p-value; CWP Low and CWP Hi=90% confidence interval for CWP; NVtxs: number of vertices in cluster; DC=diencephalon; dACC/SMA=dorsal Anterior Cingulate Cortex/Supplementary Motor Area; DMN=Default-Mode Network; EC=entorhinal cortex; R=right; PCC/RSC=posterior cingulate cortex/retrosplenial cortex.

**Supplementary Table 3.** Hippocampal (HP) functional connectivity analysis: comparison between nc-MCI and HC subjects.

| Cortical analysis |  |  |  |  |  |  |  |  |  |
| --- | --- | --- | --- | --- | --- | --- | --- | --- | --- |
| Regions included in the cluster | Size (mm<sup>2</sup>) | MNI - X | MNI - Y | MNI - Z | CWP | CWP Low | CWP Hi | NVtxs | WghtVtx |
| Seed: R-HP |  |  |  |  |  |  |  |  |  |
| L-mPFC | 692.85 | 9.6 | 55.9 | -3.7 | 0.0003 | 0 | 0.0006 | 968 | -2824.15 |
| R-mPFC | 537.65 | -12.2 | 54.6 | 6.1 | 0.00509 | 0.0036 | 0.00659 | 814 | -2255.72 |
| Sub-cortical analysis |  |  |  |  |  |  |  |  |  |
| Regions included in the cluster | Size (mm<sup>2</sup>) | MNI - X | MNI - Y | MNI - Z | GRF CWP | Cortical network associated to cerebellar region |  |  |  |
| Seed: R-HP |  |  |  |  |  |  |  |  |  |
| R-Accumbens | 16 | 10 | 7 | -13 | 0.0000052 |  |  |  |  |
| R-Cerebellum | 32 | 14 | -81 | -41 | 0.0000175 | DMN |  |  |  |
| R-Cerebellum | 32 | 18 | -75 | -55 | 0.0000349 | DAN |  |  |  |
| R-Caudate | 40 | 16 | 25 | 7 | 0.0001305 |  |  |  |  |
| L-SC | 64 | -6 | -37 | -3 | 0.0004355 |  |  |  |  |
| R-Cerebellum | 64 | 32 | -63 | -59 | 0.0008709 | DAN |  |  |  |
| L-Caudate | 152 | -14 | 23 | 7 | 0.0082139 |  |  |  |  |
| L-Thalamus | 96 | -14 | -25 | 1 | 0.0014129 |  |  |  |  |
| L-Putamen | 112 | -20 | 19 | -5 | 0.0025668 |  |  |  |  |
| Hypothalamus | 136 | 0 | 3 | -11 | 0.004481 |  |  |  |  |
| L-Hippocampus | 176 | -22 | -19 | -21 | 0.0288206 |  |  |  |  |
| Seed: L-HP |  |  |  |  |  |  |  |  |  |
| Brainstem | 1800 | 4 | -37 | -41 | 0.0214123 |  |  |  |  |

*Abbreviations:* CWP=cluster wise p-value; CWP Low and CWP Hi=90% confidence interval for CWP; NVtxs: number of vertices in cluster; L=left; mPFC=medial Prefrontal Cortex; R=right; SC=Superior Colliculus.

**Supplementary Table 4.** Hippocampal (HP) functional connectivity analysis: comparison between nc-MCI and c-MCI patients.

| *Sub-cortical analysis* |  |  |  |  |  |  |
| --- | --- | --- | --- | --- | --- | --- |
| Regions included in the cluster | Size (mm<sup>2</sup>) | MNI - X | MNI - Y | MNI - Z | GRF CWP | Cortical network associated to cerebellar region |
| **Seed: R-HP** |  |  |  |  |  |  |
| L-Cerebellum | 32 | -30 | -47 | -33 | 0.008915 | SN/CON |
| R-Cerebellum | 1736 | 38 | -75 | -41 | 0.0314906 | DMN |
| R-Cerebellum | 96 | 32 | -51 | -37 | 0.0444732 | FPN |

*Abbreviations:* CWP=cluster wise p-value; CWP Low and CWP Hi=90% confidence interval for CWP; NVtxs: number of vertices in cluster; L=left; R=right.

**Supplementary Table 5.** Entorhinal (EC) functional connectivity analysis: comparison between nc-MCI and HC subjects.

| *Sub-cortical analysis* |  |  |  |  |  |  |
| --- | --- | --- | --- | --- | --- | --- |
| Regions included in the cluster | Size (mm<sup>2</sup>) | MNI - X | MNI - Y | MNI - Z | GRF CWP | Cortical network associated to cerebellar region |
| **Seed: L-EC** |  |  |  |  |  |  |
| R-Cerebellum | 1408 | 6 | -51 | -31 | 0.0420196 | SMN, CON/SN, Limbic Network |

*Abbreviations:* CWP=cluster wise p-value; CWP Low and CWP Hi=90% confidence interval for CWP; NVtxs: number of vertices in cluster; L=left; R=right.

**Supplementary Table 6.** Entorhinal (EC) functional connectivity analysis: comparison between c-MCI and HC subjects.

| Cortical analysis |  |  |  |  |  |  |  |  |  |
| --- | --- | --- | --- | --- | --- | --- | --- | --- | --- |
| Regions included in the cluster | Size (mm<sup>2</sup>) | MNI - X | MNI - Y | MNI - Z | CWP | CWP Low | CWP Hi | NVtxs | WghtVtx |
| Seed: R-EC |  |  |  |  |  |  |  |  |  |
| R-SMA/MFG | 586.24 | 21.7 | 0.9 | 51.6 | 0.0003 | 0 | 0.0006 | 1330 | 3668.35 |
| Lat. Occipital | 891.69 | -28.8 | -68 | 23 | 0.0003 | 0 | 0.0006 | 1556 | 4319.27 |
| R-SPL | 540.58 | -32 | -53.1 | 54.8 | 0.0009 | 0.0003 | 0.0015 | 1375 | 3669.54 |
| Sub-cortical analysis |  |  |  |  |  |  |  |  |  |
| Regions included in the cluster | Size (mm<sup>2</sup>) | MNI - X | MNI - Y | MNI - Z | GRF CWP | Cortical network associated to cerebellar region |  |  |  |
| Seed: R-EC |  |  |  |  |  |  |  |  |  |
| L-Putamen | 40 | -32 | -5 | -13 | 0 |  |  |  |  |
| L-Thalamus | 16 | -4 | -9 | 1 | 0 |  |  |  |  |
| R-Hypothalamus | 56 | 12 | 5 | -15 | 0.0000002 |  |  |  |  |
| L-Cerebellum | 56 | -18 | -61 | -27 | 0.0000002 | Limbic Network |  |  |  |
| L-Cerebellum | 56 | -10 | -81 | -41 | 0.0000009 | DMN |  |  |  |
| L-Cerebellum | 64 | -16 | -45 | -45 | 0.0000036 | DAN/FPN |  |  |  |
| L-Pallidum | 72 | -16 | 3 | -5 | 0.0000138 |  |  |  |  |
| R-Thalamus | 80 | 28 | -33 | -1 | 0.0000517 |  |  |  |  |
| L- Thalamus | 88 | -16 | -23 | 3 | 0.0000978 |  |  |  |  |
| L-Cerebellum | 96 | -14 | -87 | -43 | 0.0001852 | DMN |  |  |  |
| L-Cerebellum | 96 | -12 | -71 | -49 | 0.0003705 | DAN |  |  |  |
| R-Cerebellum | 240 | 18 | -63 | -31 | 240 |  |  |  |  |
| Brainstem | 240 | 10 | -29 | -33 | 240 |  |  |  |  |
| Brainstem | 1576 | 2 | -43 | -33 | 1576 |  |  |  |  |
| R-Pallidum | 296 | 20 | 1 | -3 | 296 |  |  |  |  |

*Abbreviations:* CWP=cluster wise p-value; CWP Low and CWP Hi=90% confidence interval for CWP; NVtxs: number of vertices in cluster; L=left; R=right; SMA/MFG=Supplementary Motor Area/Middle Frontal Gyrus; SPL=Superior Parietal Lobe.

**Supplementary Table 7.**
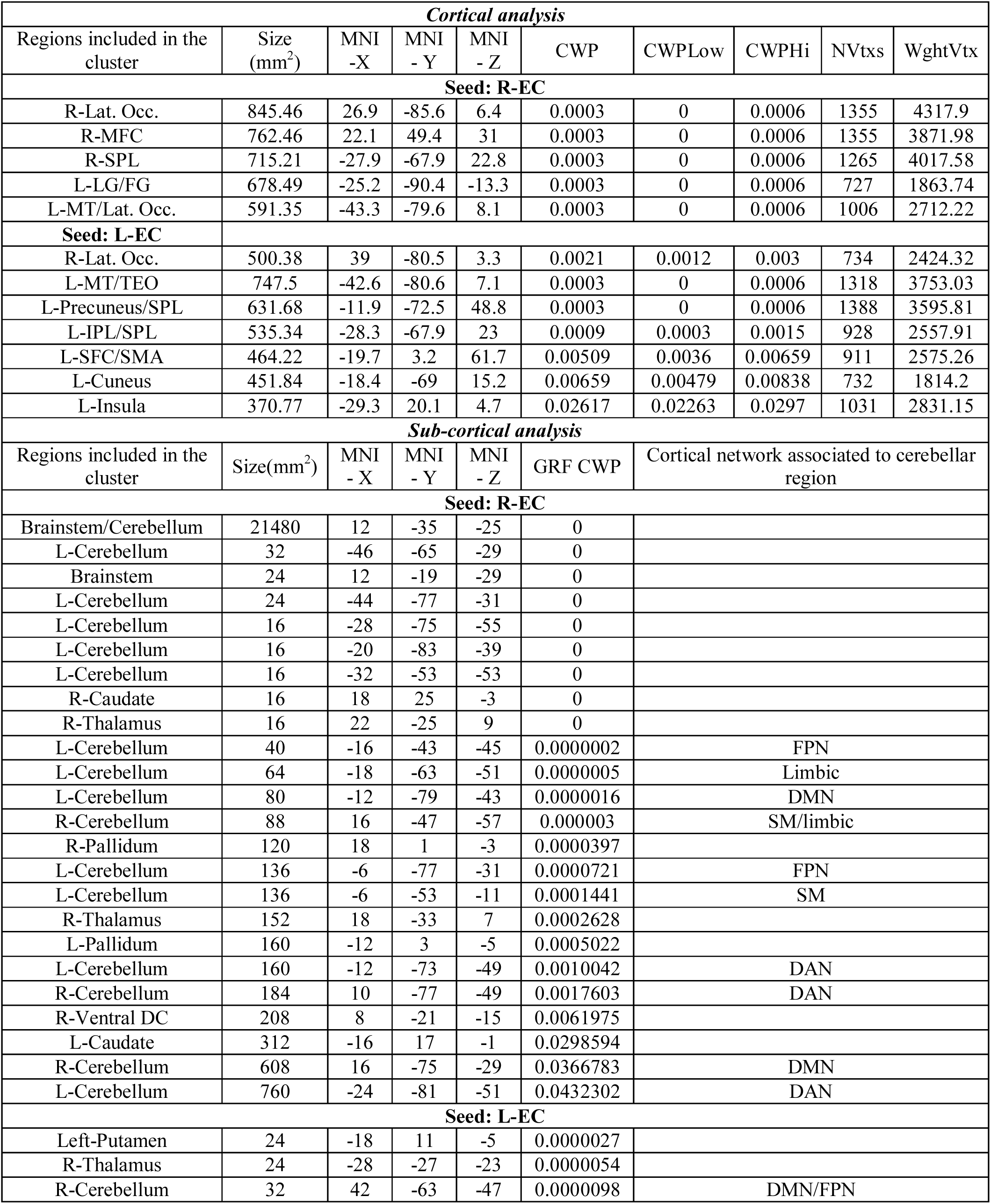
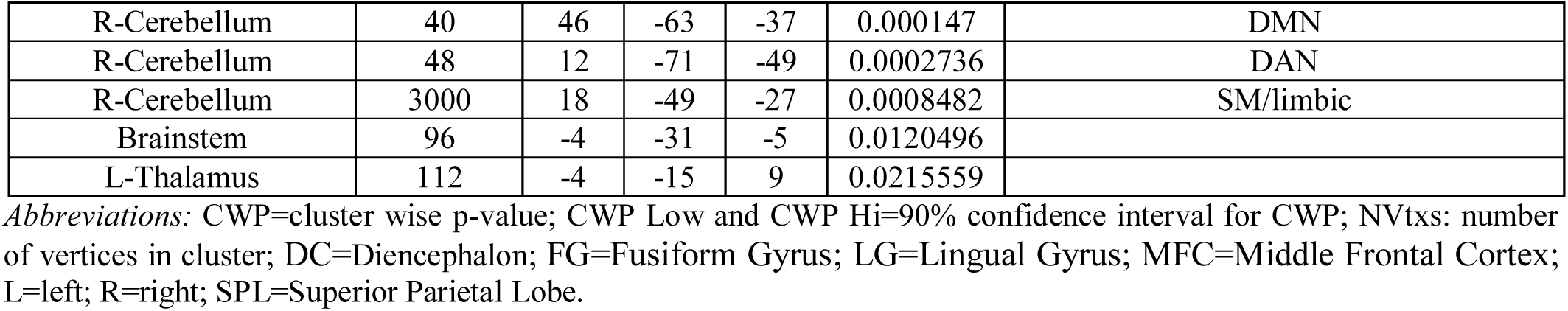
Entorhinal (EC) functional connectivity analysis: comparison between nc-MCI and c-MCI patients.

| Cortical analysis |  |  |  |  |  |  |  |  |  |
| --- | --- | --- | --- | --- | --- | --- | --- | --- | --- |
| Regions included in the cluster | Size (mm<sup>2</sup>) | MNI -X | MNI - Y | MNI - Z | CWP | CWPLow | CWPHi | NVtxs | WghtVtx |
| Seed: R-EC |  |  |  |  |  |  |  |  |  |
| R-Lat. Occ. | 845.46 | 26.9 | -85.6 | 6.4 | 0.0003 | 0 | 0.0006 | 1355 | 4317.9 |
| R-MFC | 762.46 | 22.1 | 49.4 | 31 | 0.0003 | 0 | 0.0006 | 1355 | 3871.98 |
| R-SPL | 715.21 | -27.9 | -67.9 | 22.8 | 0.0003 | 0 | 0.0006 | 1265 | 4017.58 |
| L-LG/FG | 678.49 | -25.2 | -90.4 | -13.3 | 0.0003 | 0 | 0.0006 | 727 | 1863.74 |
| L-MT/Lat. Occ. | 591.35 | -43.3 | -79.6 | 8.1 | 0.0003 | 0 | 0.0006 | 1006 | 2712.22 |
| Seed: L-EC |  |  |  |  |  |  |  |  |  |
| R-Lat. Occ. | 500.38 | 39 | -80.5 | 3.3 | 0.0021 | 0.0012 | 0.003 | 734 | 2424.32 |
| L-MT/TEO | 747.5 | -42.6 | -80.6 | 7.1 | 0.0003 | 0 | 0.0006 | 1318 | 3753.03 |
| L-Precuneus/SPL | 631.68 | -11.9 | -72.5 | 48.8 | 0.0003 | 0 | 0.0006 | 1388 | 3595.81 |
| L-IPL/SPL | 535.34 | -28.3 | -67.9 | 23 | 0.0009 | 0.0003 | 0.0015 | 928 | 2557.91 |
| L-SFC/SMA | 464.22 | -19.7 | 3.2 | 61.7 | 0.00509 | 0.0036 | 0.00659 | 911 | 2575.26 |
| L-Cuneus | 451.84 | -18.4 | -69 | 15.2 | 0.00659 | 0.00479 | 0.00838 | 732 | 1814.2 |
| L-Insula | 370.77 | -29.3 | 20.1 | 4.7 | 0.02617 | 0.02263 | 0.0297 | 1031 | 2831.15 |
| Sub-cortical analysis |  |  |  |  |  |  |  |  |  |
| Regions included in the cluster | Size(mm<sup>2</sup>) | MNI - X | MNI - Y | MNI - Z | GRF CWP | Cortical network associated to cerebellar region |  |  |  |
| Seed: R-EC |  |  |  |  |  |  |  |  |  |
| Brainstem/Cerebellum | 21480 | 12 | -35 | -25 | 0 |  |  |  |  |
| L-Cerebellum | 32 | -46 | -65 | -29 | 0 |  |  |  |  |
| Brainstem | 24 | 12 | -19 | -29 | 0 |  |  |  |  |
| L-Cerebellum | 24 | -44 | -77 | -31 | 0 |  |  |  |  |
| L-Cerebellum | 16 | -28 | -75 | -55 | 0 |  |  |  |  |
| L-Cerebellum | 16 | -20 | -83 | -39 | 0 |  |  |  |  |
| L-Cerebellum | 16 | -32 | -53 | -53 | 0 |  |  |  |  |
| R-Caudate | 16 | 18 | 25 | -3 | 0 |  |  |  |  |
| R-Thalamus | 16 | 22 | -25 | 9 | 0 |  |  |  |  |
| L-Cerebellum | 40 | -16 | -43 | -45 | 0.0000002 | FPN |  |  |  |
| L-Cerebellum | 64 | -18 | -63 | -51 | 0.0000005 | Limbic |  |  |  |
| L-Cerebellum | 80 | -12 | -79 | -43 | 0.0000016 | DMN |  |  |  |
| R-Cerebellum | 88 | 16 | -47 | -57 | 0.000003 | SM/limbic |  |  |  |
| R-Pallidum | 120 | 18 | 1 | -3 | 0.0000397 |  |  |  |  |
| L-Cerebellum | 136 | -6 | -77 | -31 | 0.0000721 | FPN |  |  |  |
| L-Cerebellum | 136 | -6 | -53 | -11 | 0.0001441 | SM |  |  |  |
| R-Thalamus | 152 | 18 | -33 | 7 | 0.0002628 |  |  |  |  |
| L-Pallidum | 160 | -12 | 3 | -5 | 0.0005022 |  |  |  |  |
| L-Cerebellum | 160 | -12 | -73 | -49 | 0.0010042 | DAN |  |  |  |
| R-Cerebellum | 184 | 10 | -77 | -49 | 0.0017603 | DAN |  |  |  |
| R-Ventral DC | 208 | 8 | -21 | -15 | 0.0061975 |  |  |  |  |
| L-Caudate | 312 | -16 | 17 | -1 | 0.0298594 |  |  |  |  |
| R-Cerebellum | 608 | 16 | -75 | -29 | 0.0366783 | DMN |  |  |  |
| L-Cerebellum | 760 | -24 | -81 | -51 | 0.0432302 | DAN |  |  |  |
| Seed: L-EC |  |  |  |  |  |  |  |  |  |
| Left-Putamen | 24 | -18 | 11 | -5 | 0.0000027 |  |  |  |  |
| R-Thalamus | 24 | -28 | -27 | -23 | 0.0000054 |  |  |  |  |
| R-Cerebellum | 32 | 42 | -63 | -47 | 0.0000098 | DMN/FPN |  |  |  |

|  |  |  |  |  |  |  |
| --- | --- | --- | --- | --- | --- | --- |
| R-Cerebellum | 40 | 46 | -63 | -37 | 0.000147 | DMN |
| R-Cerebellum | 48 | 12 | -71 | -49 | 0.0002736 | DAN |
| R-Cerebellum | 3000 | 18 | -49 | -27 | 0.0008482 | SM/limbic |
| Brainstem | 96 | -4 | -31 | -5 | 0.0120496 |  |
| L-Thalamus | 112 | -4 | -15 | 9 | 0.0215559 |  |

*Abbreviations:* CWP=cluster wise p-value; CWP Low and CWP Hi=90% confidence interval for CWP; NVtxs: number of vertices in cluster; DC=Diencephalon; FG=Fusiform Gyrus; LG=Lingual Gyrus; MFC=Middle Frontal Cortex; L=left; R=right; SPL=Superior Parietal Lobe.

**Supplementary Table 8.**
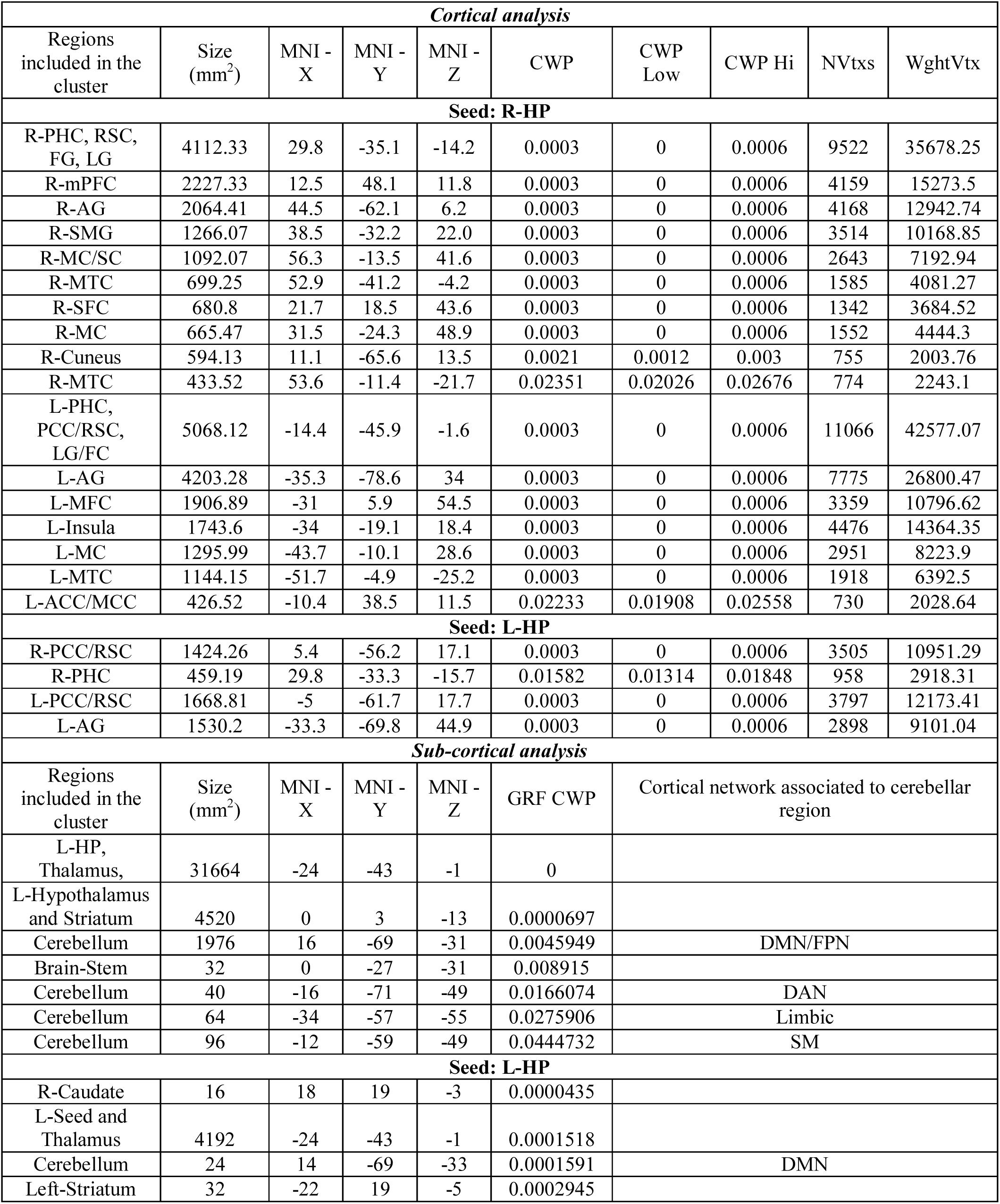
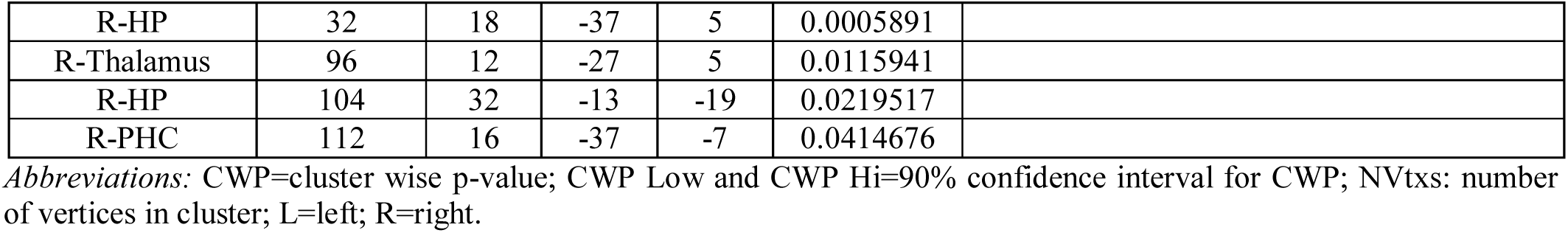
Hippocampal (HP) functional connectivity analysis: comparison between AD and nc-MCI patients.

| Cortical analysis |  |  |  |  |  |  |  |  |  |
| --- | --- | --- | --- | --- | --- | --- | --- | --- | --- |
| Regions included in the cluster | Size (mm<sup>2</sup>) | MNI - X | MNI - Y | MNI - Z | CWP | CWP Low | CWP Hi | NVtxs | WghtVtx |
| Seed: R-HP |  |  |  |  |  |  |  |  |  |
| R-PHC, RSC, FG, LG | 4112.33 | 29.8 | -35.1 | -14.2 | 0.0003 | 0 | 0.0006 | 9522 | 35678.25 |
| R-mPFC | 2227.33 | 12.5 | 48.1 | 11.8 | 0.0003 | 0 | 0.0006 | 4159 | 15273.5 |
| R-AG | 2064.41 | 44.5 | -62.1 | 6.2 | 0.0003 | 0 | 0.0006 | 4168 | 12942.74 |
| R-SMG | 1266.07 | 38.5 | -32.2 | 22.0 | 0.0003 | 0 | 0.0006 | 3514 | 10168.85 |
| R-MC/SC | 1092.07 | 56.3 | -13.5 | 41.6 | 0.0003 | 0 | 0.0006 | 2643 | 7192.94 |
| R-MTC | 699.25 | 52.9 | -41.2 | -4.2 | 0.0003 | 0 | 0.0006 | 1585 | 4081.27 |
| R-SFC | 680.8 | 21.7 | 18.5 | 43.6 | 0.0003 | 0 | 0.0006 | 1342 | 3684.52 |
| R-MC | 665.47 | 31.5 | -24.3 | 48.9 | 0.0003 | 0 | 0.0006 | 1552 | 4444.3 |
| R-Cuneus | 594.13 | 11.1 | -65.6 | 13.5 | 0.0021 | 0.0012 | 0.003 | 755 | 2003.76 |
| R-MTC | 433.52 | 53.6 | -11.4 | -21.7 | 0.02351 | 0.02026 | 0.02676 | 774 | 2243.1 |
| L-PHC, PCC/RSC, LG/FC | 5068.12 | -14.4 | -45.9 | -1.6 | 0.0003 | 0 | 0.0006 | 11066 | 42577.07 |
| L-AG | 4203.28 | -35.3 | -78.6 | 34 | 0.0003 | 0 | 0.0006 | 7775 | 26800.47 |
| L-MFC | 1906.89 | -31 | 5.9 | 54.5 | 0.0003 | 0 | 0.0006 | 3359 | 10796.62 |
| L-Insula | 1743.6 | -34 | -19.1 | 18.4 | 0.0003 | 0 | 0.0006 | 4476 | 14364.35 |
| L-MC | 1295.99 | -43.7 | -10.1 | 28.6 | 0.0003 | 0 | 0.0006 | 2951 | 8223.9 |
| L-MTC | 1144.15 | -51.7 | -4.9 | -25.2 | 0.0003 | 0 | 0.0006 | 1918 | 6392.5 |
| L-ACC/MCC | 426.52 | -10.4 | 38.5 | 11.5 | 0.02233 | 0.01908 | 0.02558 | 730 | 2028.64 |
| Seed: L-HP |  |  |  |  |  |  |  |  |  |
| R-PCC/RSC | 1424.26 | 5.4 | -56.2 | 17.1 | 0.0003 | 0 | 0.0006 | 3505 | 10951.29 |
| R-PHC | 459.19 | 29.8 | -33.3 | -15.7 | 0.01582 | 0.01314 | 0.01848 | 958 | 2918.31 |
| L-PCC/RSC | 1668.81 | -5 | -61.7 | 17.7 | 0.0003 | 0 | 0.0006 | 3797 | 12173.41 |
| L-AG | 1530.2 | -33.3 | -69.8 | 44.9 | 0.0003 | 0 | 0.0006 | 2898 | 9101.04 |
| Sub-cortical analysis |  |  |  |  |  |  |  |  |  |
| Regions included in the cluster | Size (mm<sup>2</sup>) | MNI - X | MNI - Y | MNI - Z | GRF CWP | Cortical network associated to cerebellar region |  |  |  |
| L-HP, Thalamus, | 31664 | -24 | -43 | -1 | 0 |  |  |  |  |
| L-Hypothalamus and Striatum | 4520 | 0 | 3 | -13 | 0.0000697 |  |  |  |  |
| Cerebellum | 1976 | 16 | -69 | -31 | 0.0045949 | DMN/FPN |  |  |  |
| Brain-Stem | 32 | 0 | -27 | -31 | 0.008915 |  |  |  |  |
| Cerebellum | 40 | -16 | -71 | -49 | 0.0166074 | DAN |  |  |  |
| Cerebellum | 64 | -34 | -57 | -55 | 0.0275906 | Limbic |  |  |  |
| Cerebellum | 96 | -12 | -59 | -49 | 0.0444732 | SM |  |  |  |
| Seed: L-HP |  |  |  |  |  |  |  |  |  |
| R-Caudate | 16 | 18 | 19 | -3 | 0.0000435 |  |  |  |  |
| L-Seed and Thalamus | 4192 | -24 | -43 | -1 | 0.0001518 |  |  |  |  |
| Cerebellum | 24 | 14 | -69 | -33 | 0.0001591 | DMN |  |  |  |
| Left-Striatum | 32 | -22 | 19 | -5 | 0.0002945 |  |  |  |  |

|  |  |  |  |  |  |
| --- | --- | --- | --- | --- | --- |
| R-HP | 32 | 18 | -37 | 5 | 0.0005891 |
| R-Thalamus | 96 | 12 | -27 | 5 | 0.0115941 |
| R-HP | 104 | 32 | -13 | -19 | 0.0219517 |
| R-PHC | 112 | 16 | -37 | -7 | 0.0414676 |

*Abbreviations:* CWP=cluster wise p-value; CWP Low and CWP Hi=90% confidence interval for CWP; NVtxs: number of vertices in cluster; L=left; R=right.

**Supplementary Table 9.** Entorhinal (EC) functional connectivity analysis: comparison between AD and c-MCI patients.

| *Sub-cortical analysis* |  |  |  |  |  |
| --- | --- | --- | --- | --- | --- |
| Regions included in the cluster | Size (mm<sup>2</sup>) | MNI - X | MNI - Y | MNI - Z | GRF CWP |
| **Seed: R-EC** |  |  |  |  |  |
| Brainstem | 1280 | -4 | -43 | -53 | 0.0356449 |
| R-Amygdala | 2680 | 18 | -13 | -23 | 0.003215 |

*Abbreviations:* CWP=cluster wise p-value; CWP Low and CWP Hi=90% confidence interval for CWP; NVtxs: number of vertices in cluster; L=left; R=right.

**Supplementary Table 10.** Whole brain correlation analysis between hippocampal (HP) functional connectivity and CSF levels of Aβ_1–42_, p-tau_181_ and p-tau_181_/A_β1–42_.

|  | Size(mm<sup>2</sup>) | MNI<br>- X | MNI<br>- Y | MNI<br>- Z | CWP | CWP Low | CWP Hi | NVtxs | WghtVtx |
| --- | --- | --- | --- | --- | --- | --- | --- | --- | --- |
| R-HP FC -Whole brain correlation with Aβ<sub>1-42</sub> |  |  |  |  |  |  |  |  |  |
| R-mPFC | 786.42 | 10.8 | 63.6 | 13.6 | 0.0003 | 0 | 0.0006 | 1147 | 3120.78 |
| R-PCC/RSC | 549.84 | 5.7 | -54.1 | 12.7 | 0.003 | 0.0018 | 0.00419 | 1311 | 3711.62 |
| R-Precuneus | 496.81 | 9.1 | -60.1 | 40.5 | 0.00867 | 0.00659 | 0.01076 | 1251 | 3808.67 |
| L-mPFC | 604.42 | -7.2 | 49.8 | 39.5 | 0.0012 | 0.0006 | 0.0021 | 995 | 2684.29 |
| L-SFC/MFC | 559.88 | -18.7 | 31.1 | 39.3 | 0.0033 | 0.0021 | 0.00449 | 1054 | 3030.96 |
| R-HP FC - Whole brain correlation with p-tau<sub>181</sub> |  |  |  |  |  |  |  |  |  |
| R-Precuneus | 1046.32 | 8.7 | -60.7 | 40.2 | 0.0003 | 0 | 0.0006 | 2299 | -7598.39 |
| R-PCC/RSC | 917.89 | 6 | -48.7 | 17.4 | 0.0003 | 0 | 0.0006 | 2298 | -7649.66 |
| R-AG | 768.87 | 38.7 | -73.3 | 29.7 | 0.0003 | 0 | 0.0006 | 1303 | -3468.07 |
| R-SFC/MFC | 672.05 | 26 | 28.1 | 35.4 | 0.0003 | 0 | 0.0006 | 1266 | -3260.6 |
| L-PCC/RSC | 2832.51 | -4 | -22.7 | 32.7 | 0.0003 | 0 | 0.0006 | 6123 | -19030.07 |
| L-AG | 2049.72 | -43.7 | -73.7 | 27.5 | 0.0003 | 0 | 0.0006 | 4084 | -11869.84 |
| L-SFC/MFC | 865.24 | -27.8 | 28.9 | 35.5 | 0.0003 | 0 | 0.0006 | 1422 | -3894.61 |
| R-HP FC - Whole brain correlation with p-tau<sub>181</sub>/Aβ<sub>1-42</sub> |  |  |  |  |  |  |  |  |  |
| R-Precuneus | 2815.23 | 7.2 | -66.8 | 41.8 | 0.0003 | 0 | 0.0006 | 6643 | -22776.81 |
| R-SFC/MFC | 1465.37 | 26 | 28.1 | 35.4 | 0.0003 | 0 | 0.0006 | 2632 | -7204.94 |
| R-AG | 1245.97 | 47.3 | -57.4 | 38.8 | 0.0003 | 0 | 0.0006 | 2279 | -6822.83 |
| R-mPFC | 1033.64 | 12.4 | 44.5 | 16.4 | 0.0003 | 0 | 0.0006 | 1505 | -4364.17 |
| L-PCC/RSC | 3383.28 | -4.1 | -23.2 | 32.4 | 0.0003 | 0 | 0.0006 | 7463 | -24119.01 |
| L-AG | 2207.16 | -43.7 | -73.7 | 27.5 | 0.0003 | 0 | 0.0006 | 4378 | -13428.81 |
| L-SFC/MFC | 1268.59 | -26.7 | 11.8 | 43.2 | 0.0003 | 0 | 0.0006 | 2188 | -6465.53 |
| L-Cuneus | 505.87 | -14 | -94.7 | -6.9 | 0.00778 | 0.00599 | 0.00987 | 679 | -1920.19 |
|  |  |  |  |  | GRF CWP | Cortical Network associated to cerebellar region |  |  |  |
| R-Cerebellum | 16 | 16 | -77 | -19 | 0.0001493 | DMN |  |  |  |
| L-Cerebellum | 88 | -50 | -71 | -39 | 0.0491381 | Limbic Network |  |  |  |
| L-Cerebellum | 40 | -32 | -47 | -31 | 0.0002782 | DMN |  |  |  |
| Brainstem | 192 | -28 | -23 | 11 | 0.0067947 |  |  |  |  |
| R-Putamen | 80 | 34 | -1 | -9 | 0.0261405 |  |  |  |  |
| R-Thalamus | 64 | 16 | -33 | -1 | 0.0073613 |  |  |  |  |
| R-Caudate | 40 | 10 | -1 | 15 | 0.0005565 |  |  |  |  |
| R-Accumbens | 176 | 10 | 11 | -9 | 0.0017803 |  |  |  |  |
| R-Amygdala | 248 | 22 | -7 | -21 | 0.0461488 |  |  |  |  |
| L-HP FC - Whole brain correlation with p-tau<sub>181</sub>/Aβ<sub>1-42</sub> |  |  |  |  |  |  |  |  |  |
| L-Cerebellum | 80 | -46 | -39 | -9 | 0.0390553 | DMN |  |  |  |

*Abbreviations:* CWP=cluster wise p-value; CWP Low and CWP Hi= 90% confidence interval for CWP; NVtxs: number of vertices in cluster; DC=Diencephalon; FG=Fusiform Gyrus; LG=Lingual Gyrus; MFC=Middle Frontal Cortex; L=left; R=right; SPL=Superior Parietal Lobe.

## REFERENCES

Apostolova LG, Dutton RA, Dinov ID, Hayashi KM, Toga AW, Cummings JL, Thompson PM. 2006. Conversion of mild cognitive impairment to Alzheimer disease predicted by hippocampal atrophy maps. Archives of neurology 63:693–699.

Badhwar A, Tam A, Dansereau C, Orban P, Hoffstaedter F, Bellec P. 2017. Resting-state network dysfunction in Alzheimer’s disease: A systematic review and meta-analysis. Alzheimer’s & dementia 8:73–85.

Barker GR, Bird F, Alexander V, Warburton EC. 2007. Recognition memory for objects, place, and temporal order: a disconnection analysis of the role of the medial prefrontal cortex and perirhinal cortex. J Neurosci 27:2948–2957.

Bondi MW, Houston WS, Eyler LT, Brown GG. 2005. fMRI evidence of compensatory mechanisms in older adults at genetic risk for Alzheimer disease. Neurology 64:501–508.

Bookheimer SY, Strojwas MH, Cohen MS, Saunders AM, Pericak-Vance MA, Mazziotta JC, Small GW. 2000. Patterns of brain activation in people at risk for Alzheimer’s disease. The New England journal of medicine 343:450–456.

Braak E, Braak H, Mandelkow EM. 1994. A sequence of cytoskeleton changes related to the formation of neurofibrillary tangles and neuropil threads. Acta neuropathologica 87:554–567.

Braak H, Feldengut S, Del Tredici K. 2013. [Pathogenesis and prevention of Alzheimer’s disease: when and in what way does the pathological process begin?]. Der Nervenarzt 84:477–482.

Buckner RL, Krienen FM, Castellanos A, Diaz JC, Yeo BT. 2011. The organization of the human cerebellum estimated by intrinsic functional connectivity. Journal of neurophysiology 106:2322–2345.

Buckner RL, Snyder AZ, Shannon BJ, LaRossa G, Sachs R, Fotenos AF, Sheline YI, Klunk WE, Mathis CA, Morris JC, Mintun MA. 2005. Molecular, structural, and functional characterization of Alzheimer’s disease: evidence for a relationship between default activity, amyloid, and memory. J Neurosci 25:7709–7717.

Busse A, Hensel A, Guhne U, Angermeyer MC, Riedel-Heller SG. 2006. Mild cognitive impairment: long-term course of four clinical subtypes. Neurology 67:2176–2185.

Cabeza R, Grady CL, Nyberg L, McIntosh AR, Tulving E, Kapur S, Jennings JM, Houle S, Craik FI. 1997. Age-related differences in neural activity during memory encoding and retrieval: a positron emission tomography study. J Neurosci 17:391–400.

Canto CB, Wouterlood FG, Witter MP. 2008. What does the anatomical organization of the entorhinal cortex tell us? Neural Plast 2008:381243.

Canuet L, Pusil S, Lopez ME, Bajo R, Pineda-Pardo JA, Cuesta P, Galvez G, Gaztelu JM, Lourido D, Garcia-Ribas G, Maestu F. 2015. Network Disruption and Cerebrospinal Fluid Amyloid-Beta and Phospho-Tau Levels in Mild Cognitive Impairment. J Neurosci 35:10325–10330.

Cope TE, Rittman T, Borchert RJ, Jones PS, Vatansever D, Allinson K, Passamonti L, Vazquez Rodriguez P, Bevan-Jones WR, O’Brien JT, Rowe JB. 2018. Tau burden and the functional connectome in Alzheimer’s disease and progressive supranuclear palsy. Brain 141:550–567.

Das SR, Pluta J, Mancuso L, Kliot D, Orozco S, Dickerson BC, Yushkevich PA, Wolk DA. 2013. Increased functional connectivity within medial temporal lobe in mild cognitive impairment. Hippocampus 23:1–6.

Davis SW, Dennis NA, Daselaar SM, Fleck MS, Cabeza R. 2008. Que PASA? The posterior-anterior shift in aging. Cereb Cortex 18:1201–1209.

Desikan RS, Segonne F, Fischl B, Quinn BT, Dickerson BC, Blacker D, Buckner RL, Dale AM, Maguire RP, Hyman BT, Albert MS, Killiany RJ. 2006. An automated labeling system for subdividing the human cerebral cortex on MRI scans into gyral based regions of interest. Neuroimage 31:968–980.

Drzezga A, Becker JA, Van Dijk KR, Sreenivasan A, Talukdar T, Sullivan C, Schultz AP, Sepulcre J, Putcha D, Greve D, Johnson KA, Sperling RA. 2011. Neuronal dysfunction and disconnection of cortical hubs in non-demented subjects with elevated amyloid burden. Brain 134:1635–1646.

Eichenbaum H. 2017. Memory: Organization and Control. Annual review of psychology 68:19–45.

Elman JA, Madison CM, Baker SL, Vogel JW, Marks SM, Crowley S, O’Neil JP, Jagust WJ. 2016. Effects of Beta-Amyloid on Resting State Functional Connectivity Within and Between Networks Reflect Known Patterns of Regional Vulnerability. Cereb Cortex 26:695–707.

Fabiani M. 2012. It was the best of times, it was the worst of times: a psychophysiologist’s view of cognitive aging. Psychophysiology 49:283–304.

Ferraris M, Ghestem A, Vicente AF, Nallet-Khosrofian L, Bernard C, Quilichini PP. 2018. The Nucleus Reuniens Controls Long-Range Hippocampo-Prefrontal Gamma Synchronization during Slow Oscillations. J Neurosci 38:3026–3038.

Folstein MF, Folstein SE, McHugh PR. 1975. “Mini-mental state”. A practical method for grading the cognitive state of patients for the clinician. Journal of psychiatric research 12:189–198.

Foster CM, Kennedy KM, Horn MM, Hoagey DA, Rodrigue KM. 2018. Both hyper- and hypo-activation to cognitive challenge are associated with increased beta-amyloid deposition in healthy aging: A nonlinear effect. Neuroimage 166:285–292.

Frisoni GB, Jessen F. 2018. One step towards dementia prevention. Lancet Neurol 17:294–295.

Gilani AM, Knowles TG, Nicol CJ. 2014. Factors affecting ranging behaviour in young and adult laying hens. British poultry science 55:127–135.

Gomez-Isla T, Price JL, McKeel DW, Jr., Morris JC, Growdon JH, Hyman BT. 1996. Profound loss of layer II entorhinal cortex neurons occurs in very mild Alzheimer’s disease. J Neurosci 16:4491–4500.

Grothe MJ, Teipel SJ, Alzheimer’s Disease Neuroimaging I. 2016. Spatial patterns of atrophy, hypometabolism, and amyloid deposition in Alzheimer’s disease correspond to dissociable functional brain networks. Human brain mapping 37:35–53.

Grundman M, Sencakova D, Jack CR, Jr., Petersen RC, Kim HT, Schultz A, Weiner MF, DeCarli C, DeKosky ST, van Dyck C, Thomas RG, Thal LJ, Alzheimer’s Disease Cooperative S. 2002. Brain MRI hippocampal volume and prediction of clinical status in a mild cognitive impairment trial. Journal of molecular neuroscience: MN 19:23–27.

Hamalainen A, Tervo S, Grau-Olivares M, Niskanen E, Pennanen C, Huuskonen J, Kivipelto M, Hanninen T, Tapiola M, Vanhanen M, Hallikainen M, Helkala EL, Nissinen A, Vanninen R, Soininen H. 2007. Voxel-based morphometry to detect brain atrophy in progressive mild cognitive impairment. Neuroimage 37:1122–1131.

Hagler DJ Jr, Saygin AP, Sereno MI. 2006. Smoothing and cluster thresholding for cortical surface-based group analysis of fMRI data. Neuroimage 33: 1093–1103.

Hellyer PJ, Shanahan M, Scott G, Wise RJ, Sharp DJ, Leech R. 2014. The control of global brain dynamics: opposing actions of frontoparietal control and default mode networks on attention. J Neurosci 34:451–461.

Henneman WJ, Sluimer JD, Barnes J, van der Flier WM, Sluimer IC, Fox NC, Scheltens P, Vrenken H, Barkhof F. 2009. Hippocampal atrophy rates in Alzheimer disease: added value over whole brain volume measures. Neurology 72:999–1007.

Hillary FG, Grafman JH. 2017. Injured Brains and Adaptive Networks: The Benefits and Costs of Hyperconnectivity. Trends in cognitive sciences 21:385–401.

Hoenig MC, Bischof GN, Seemiller J, Hammes J, Kukolja J, Onur OA, Jessen F, Fliessbach K, Neumaier B, Fink GR, van Eimeren T, Drzezga A. 2018. Networks of tau distribution in Alzheimer’s disease. Brain 141:568–581.

Huijbers W, Mormino EC, Schultz AP, Wigman S, Ward AM, Larvie M, Amariglio RE, Marshall GA, Rentz DM, Johnson KA, Sperling RA. Amyloid-beta deposition in mild cognitive impairment is associated with increased hippocampal activity, atrophy and clinical progression. Brain 2015; 138(Pt 4): 1023–35.

Jack CR, Jr., Shiung MM, Gunter JL, O’Brien PC, Weigand SD, Knopman DS, Boeve BF, Ivnik RJ, Smith GE, Cha RH, Tangalos EG, Petersen RC. 2004. Comparison of different MRI brain atrophy rate measures with clinical disease progression in AD. Neurology 62:591–600.

Jones DT, Knopman DS, Gunter JL, Graff-Radford J, Vemuri P, Boeve BF, Petersen RC, Weiner MW, Jack CR, Jr., Alzheimer’s Disease Neuroimaging I. 2016. Cascading network failure across the Alzheimer’s disease spectrum. Brain 139:547–562.

Kahn I, Andrews-Hanna JR, Vincent JL, Snyder AZ, Buckner RL. 2008. Distinct cortical anatomy linked to subregions of the medial temporal lobe revealed by intrinsic functional connectivity. Journal of neurophysiology 100:129–139.

Kaplan E GH, Weintraub S, editors. 1983. The Boston Naming Test. In: Philadelphia, PA. Lea & Febiger.

Kircher TT, Weis S, Freymann K, Erb M, Jessen F, Grodd W, Heun R, Leube DT. 2007. Hippocampal activation in patients with mild cognitive impairment is necessary for successful memory encoding. J Neurol Neurosurg Psychiatry 78:812–818.

Koch W, Teipel S, Mueller S, Buerger K, Bokde AL, Hampel H, Coates U, Reiser M, Meindl T. 2010. Effects of aging on default mode network activity in resting state fMRI: does the method of analysis matter? Neuroimage 51:280–287.

Lacy JW, Stark CE. 2012. Intrinsic functional connectivity of the human medial temporal lobe suggests a distinction between adjacent MTL cortices and hippocampus. Hippocampus 22:2290–2302.

Libby LA, Ekstrom AD, Ragland JD, Ranganath C. 2012. Differential connectivity of perirhinal and parahippocampal cortices within human hippocampal subregions revealed by high-resolution functional imaging. J Neurosci 32:6550–6560.

Mitchell AJ, Shiri-Feshki M. 2009. Rate of progression of mild cognitive impairment to dementia–meta-analysis of 41 robust inception cohort studies. Acta psychiatrica Scandinavica 119:252–265.

Mohs RC, Cohen L. 1988. Alzheimer’s Disease Assessment Scale (ADAS). Psychopharmacology bulletin 24:627–628.

Mohs RC, Knopman D, Petersen RC, Ferris SH, Ernesto C, Grundman M, Sano M, Bieliauskas L, Geldmacher D, Clark C, Thal LJ. 1997. Development of cognitive instruments for use in clinical trials of antidementia drugs: additions to the Alzheimer’s Disease Assessment Scale that broaden its scope. The Alzheimer’s Disease Cooperative Study. Alzheimer disease and associated disorders 11 Suppl 2:S13–21.

Mormino EC, Smiljic A, Hayenga AO, Onami SH, Greicius MD, Rabinovici GD, Janabi M, Baker SL, Yen IV, Madison CM, Miller BL, Jagust WJ. 2011. Relationships between beta-amyloid and functional connectivity in different components of the default mode network in aging. Cereb Cortex 21:2399–2407.

Morris JC. 1993. The Clinical Dementia Rating (CDR): current version and scoring rules. Neurology 43:2412–2414.

Morris JC, Heyman A, Mohs RC, Hughes JP, van Belle G, Fillenbaum G, Mellits ED, Clark C. 1989. The Consortium to Establish a Registry for Alzheimer’s Disease (CERAD). Part I. Clinical and neuropsychological assessment of Alzheimer’s disease. Neurology 39:1159–1165.

Mutlu J, Landeau B, Gaubert M, de La Sayette V, Desgranges B, Chetelat G. 2017. Distinct influence of specific versus global connectivity on the different Alzheimer’s disease biomarkers. Brain 140:3317–3328.

Nasreddine ZS, Phillips NA, Bedirian V, Charbonneau S, Whitehead V, Collin I, Cummings JL, Chertkow H. 2005. The Montreal Cognitive Assessment, MoCA: a brief screening tool for mild cognitive impairment. Journal of the American Geriatrics Society 53:695–699.

Oh H, Jagust WJ. 2013. Frontotemporal network connectivity during memory encoding is increased with aging and disrupted by beta-amyloid. J Neurosci 33:18425–18437.

Palmqvist S, Scholl M, Strandberg O, Mattsson N, Stomrud E, Zetterberg H, Blennow K, Landau S, Jagust W, Hansson O. 2017. Earliest accumulation of beta-amyloid occurs within the default-mode network and concurrently affects brain connectivity. Nat Commun 8:1214.

Park DC, Reuter-Lorenz P. 2009. The adaptive brain: aging and neurocognitive scaffolding. Annual review of psychology 60:173–196.

Pasquini L, Scherr M, Tahmasian M, Meng C, Myers NE, Ortner M, Muhlau M, Kurz A, Forstl H, Zimmer C, Grimmer T, Wohlschlager AM, Riedl V, Sorg C. 2015. Link between hippocampus’ raised local and eased global intrinsic connectivity in AD. Alzheimers Dement 11:475–484.

Petersen RC, Aisen PS, Beckett LA, Donohue MC, Gamst AC, Harvey DJ, Jack CR, Jr., Jagust WJ, Shaw LM, Toga AW, Trojanowski JQ, Weiner MW. 2010. Alzheimer’s Disease Neuroimaging Initiative (ADNI): clinical characterization. Neurology 74:201–209.

Pfeffer RI, Kurosaki TT, Harrah CH, Jr., Chance JM, Filos S. 1982. Measurement of functional activities in older adults in the community. Journal of gerontology 37:323–329.

Preston AR, Eichenbaum H. 2013. Interplay of hippocampus and prefrontal cortex in memory. Current biology: CB 23:R764–773.

Putcha D, Brickhouse M, O’Keefe K, Sullivan C, Rentz D, Marshall G, Dickerson B, Sperling R. 2011. Hippocampal hyperactivation associated with cortical thinning in Alzheimer’s disease signature regions in non-demented elderly adults. J Neurosci 31:17680–17688.

Ranganath C, Ritchey M. 2012. Two cortical systems for memory-guided behaviour. Nat Rev Neurosci 13:713–726.

Rey A. editor. L’examen clinique en psychologie, Presses Universitaires de France. Paris; 1964.

Reuter-Lorenz PA, Park DC. 2014. How does it STAC up? Revisiting the scaffolding theory of aging and cognition. Neuropsychology review 24:355–370.

Schmitz JM, Green CE, Hasan KM, Vincent J, Suchting R, Weaver MF, Moeller FG, Narayana PA, Cunningham KA, Dineley KT, Lane SD. 2017. PPAR-gamma agonist pioglitazone modifies craving intensity and brain white matter integrity in patients with primary cocaine use disorder: a double-blind randomized controlled pilot trial. Addiction 112:1861–1868.

Schneider-Garces NJ, Gordon BA, Brumback-Peltz CR, Shin E, Lee Y, Sutton BP, Maclin EL, Gratton G, Fabiani M. 2010. Span, CRUNCH, and beyond: working memory capacity and the aging brain. Journal of cognitive neuroscience 22:655–669.

Schobel SA, Chaudhury NH, Khan UA, Paniagua B, Styner MA, Asllani I, Inbar BP, Corcoran CM, Lieberman JA, Moore H, Small SA. 2013. Imaging patients with psychosis and a mouse model establishes a spreading pattern of hippocampal dysfunction and implicates glutamate as a driver. Neuron 78:81–93.

Schultz AP, Chhatwal JP, Hedden T, Mormino EC, Hanseeuw BJ, Sepulcre J, Huijbers W, LaPoint M, Buckley RF, Johnson KA, Sperling RA. 2017. Phases of Hyperconnectivity and Hypoconnectivity in the Default Mode and Salience Networks Track with Amyloid and Tau in Clinically Normal Individuals. J Neurosci 37:4323–4331.

Scimeca JM, Badre D. 2012. Striatal contributions to declarative memory retrieval. Neuron 75:380–392.

Sepulcre J, Sabuncu MR, Li Q, El Fakhri G, Sperling R, Johnson KA. 2017. Tau and amyloid beta proteins distinctively associate to functional network changes in the aging brain. Alzheimers Dement 13:1261–1269.

Sestieri C, Shulman GL, Corbetta M. 2017. The contribution of the human posterior parietal cortex to episodic memory. Nat Rev Neurosci 18:183–192.

Sperling RA, Laviolette PS, O’Keefe K, O’Brien J, Rentz DM, Pihlajamaki M, Marshall G, Hyman BT, Selkoe DJ, Hedden T, Buckner RL, Becker JA, Johnson KA. 2009. Amyloid deposition is associated with impaired default network function in older persons without dementia. Neuron 63:178–188.

Spreen O SE. 1998. Compendium of neuropsychological tests. In. New York: Oxford University Press.

Stoodley CJ. 2012. The cerebellum and cognition: evidence from functional imaging studies. Cerebellum 11:352–365.

Tulving E, Kapur S, Craik FI, Moscovitch M, Houle S. 1994. Hemispheric encoding/retrieval asymmetry in episodic memory: positron emission tomography findings. Proceedings of the National Academy of Sciences of the United States of America 91:2016–2020.

van den Heuvel MP, Sporns O. 2011. Rich-club organization of the human connectome. J Neurosci 31:15775–15786.

Vann SD, Aggleton JP, Maguire EA. 2009. What does the retrosplenial cortex do? Nat Rev Neurosci 10:792–802.

Wechsler D. 1987. WMS-R: Wechsler memory scale-revised. San Antonio, TX: The Psychological Corporation.

Yassa MA, Muftuler LT, Stark CE. 2010. Ultrahigh-resolution microstructural diffusion tensor imaging reveals perforant path degradation in aged humans in vivo. Proceedings of the National Academy of Sciences of the United States of America 107:12687–12691.

Yeo BT, Krienen FM, Sepulcre J, Sabuncu MR, Lashkari D, Hollinshead M, Roffman JL, Smoller JW, Zollei L, Polimeni JR, Fischl B, Liu H, Buckner RL. 2011. The organization of the human cerebral cortex estimated by intrinsic functional connectivity. J Neurophysiol 106:1125–65, 2011.

Yu W, Krook-Magnuson E. 2015. Cognitive Collaborations: Bidirectional Functional Connectivity Between the Cerebellum and the Hippocampus. Front Syst Neurosci 9:177.

